# Cardiomyograph: a robust tool for evaluating contraction of cardiomyocytes and cardiac organoids with ECG-like reports

**DOI:** 10.64898/2026.11.17.767183

**Authors:** Xun Yuan, Kaige Li, Qi An, Kai Jiang, Jingwei Wang, Dandan Wang, Yue Xu, Yaozu Xiang

## Abstract

**Background:** Cardiomyocytes are the only cells in the human body that can beat autonomously and rhythmically. Quantification of cardiomyocyte contraction is key to understand the cellular mechanisms underlying cardiac disease and drug responses. However, temporal accuracy and spatial resolution are rarely achieved together in measurements of cardiomyocyte contraction.

**Objective:** To quantify the contraction of in vitro cardiomyocyte models accurately in both time and space.

**Methods:** Here we develop an automated, open-source software (Cardiomyograph) that allows the user to process multiple videomicroscopy to obtain contractility parameters and quantitative analysis of cardiomyocyte contraction.

**Results:** Compared with current software, Cardiomyograph uses deep learning-based motion field extraction and domain-resolved beat recognition and kinematics, which can more accurately and efficiently characterize the contraction and relaxation processes of cardiomyocytes, thereby enabling precise quantitative assessment of cardiomyocyte contraction and relaxation frequency, amplitude, waveforms and intervals. The software was validated by using both primary isolated cardiomyocytes and human-induced pluripotent stem-cell-derived cardiomyocytes, treated or not with drugs or stimulus or mutation, known to affect contractility.

**Conclusion:** Therefore, Cardiomyograph provides a high-performance platform for cardiac disease phenotype, pharmacological and toxicology screening.

## Introduction

Cardiovascular diseases remain the leading cause of mortality globally^1^, with cultured cardiomyocyte models serving as pivotal tools for investigating the pathologies^2^. As a fundamental cardiac function, the contraction of cardiomyocytes is a core and widely measured parameter in in vitro experiments^2^. Cardiomyocytes exhibit rhythmic contraction and relaxation, and in multicellular cultures they contract as an electrically coupled population^3, 4^. Therefore, reliable evaluation of contractile function requires precise detection of contraction-relaxation phases and quantification of spatiotemporal coordination within electrically coupled cardiomyocyte networks. However, few existing methods or tools adequately satisfy both criteria simultaneously.

Existing characterization approaches target distinct stages of excitation-contraction coupling in cultured cardiomyocytes^3^. Patch clamp enables high-fidelity resolution of membrane currents and action potentials; however, it suffers from inherently low throughput, imposes cellular invasiveness, and —due to its single-cell recording paradigm—fails to resolve intercellular coordination^5, 6^. Microelectrode arrays can record extracellular electrical activity; however, their applications are limited by hardware costs (resulting in low throughput)^7^, and electrode sampling capabilities (leading to limited spatial resolution)^6^. Optical calcium imaging reports calcium handling across the entire field of view, but it necessitates exogenous fluorescent indicators, which interfere with native Ca^2+^ homeostasis, and critically, the contractile timing cannot be directly measured, but can only be inferred indirectly from the temporal profile of Ca^2+^ transients^8^. Traction force microscopy can estimate single-cell forces transmitted to a calibrated substrate, yet requires substrate preparation and cannot resolve coordinated interactions among multiple cells^9^.

Video-based analysis of readily acquired microscopy recordings offers a label-free, noninvasive, and high-throughput alternative^10, 11^. Existing video-based tools broadly follow four approaches: reference-frame difference, consecutive-frame difference, optical flow, and image correlation. Reference-frame difference methods take the intensity difference between each frame and a reference frame as the contraction signal^10, 12^, and any drift of the field relative to the reference adds to this signal. Consecutive-frame difference methods measure the change between consecutive frames^11^. This measure needs no reference frame but responds to image noise as well as to motion. Optical-flow methods estimate a motion vector at each pixel^13, 14^, but they lack an accurate algorithm dedicated to microscopy recordings of cells. Image-correlation methods match small windows of the image between frames and assign one displacement to each window^15, 16^. Motion that differs within a window is averaged, and windows with little texture give no reliable match. In most of these tools, the quantitative parameters are derived from a single averaged trace of motion magnitude. Temporally, this trace lacks directional information on cellular motion, and consequently, contraction and relaxation phases are delineated solely based on peak order or intensity thresholds. Such parameters typically require manual optimization for each individual recording^10, 11, 13^. Spatially, most tools aggregate signals over the entire field of view, thereby precluding quantitative assessment of regional coordination within the culture. For disease modelling and drug evaluation, these limitations mean that contraction events are mislocated in time and that regional heterogeneity in contractile behaviour is obscured. Both consequences affect the measurements on which such studies rely.

In this paper, we present an open-source software (Cardiomyograph) that reconstructs spatiotemporally resolved mechanical cardiograms of cardiomyocyte cultures from label-free microscopy videos with high fidelity and full automation. Compared with current software tools, Cardiomyograph uses deep-learning-based motion field extraction and domain-resolved beat recognition and kinematics, which can accurately identify the contractile domains of cardiomyocyte beating and characterize the contraction and relaxation processes, thereby enabling precise quantitative assessment of cardiomyocyte contraction and relaxation frequency, amplitude, waveforms, intervals and response sensitivity (at least 10-fold). The framework first estimates a dense displacement field and then identifies local contractile domains (Fig. 1). (a) Temporally, motion direction and magnitude are combined within each domain to accurately detect contraction-relaxation phases and extract contraction parameters. (b) Spatially, each contractile domain is analysed separately, so Cardiomyograph resolves focal activity obscured by whole-field averaging and quantifies the coordination among domains. Specifically, Cardiomyograph is an artificial intelligence framework offering broad utility across ultra-sensitive drug response profiling, physiologically relevant cardiovascular disease modelling, unique mechanism discovery, and identification of abnormal waveform phenotypes.

**Fig. 1.**
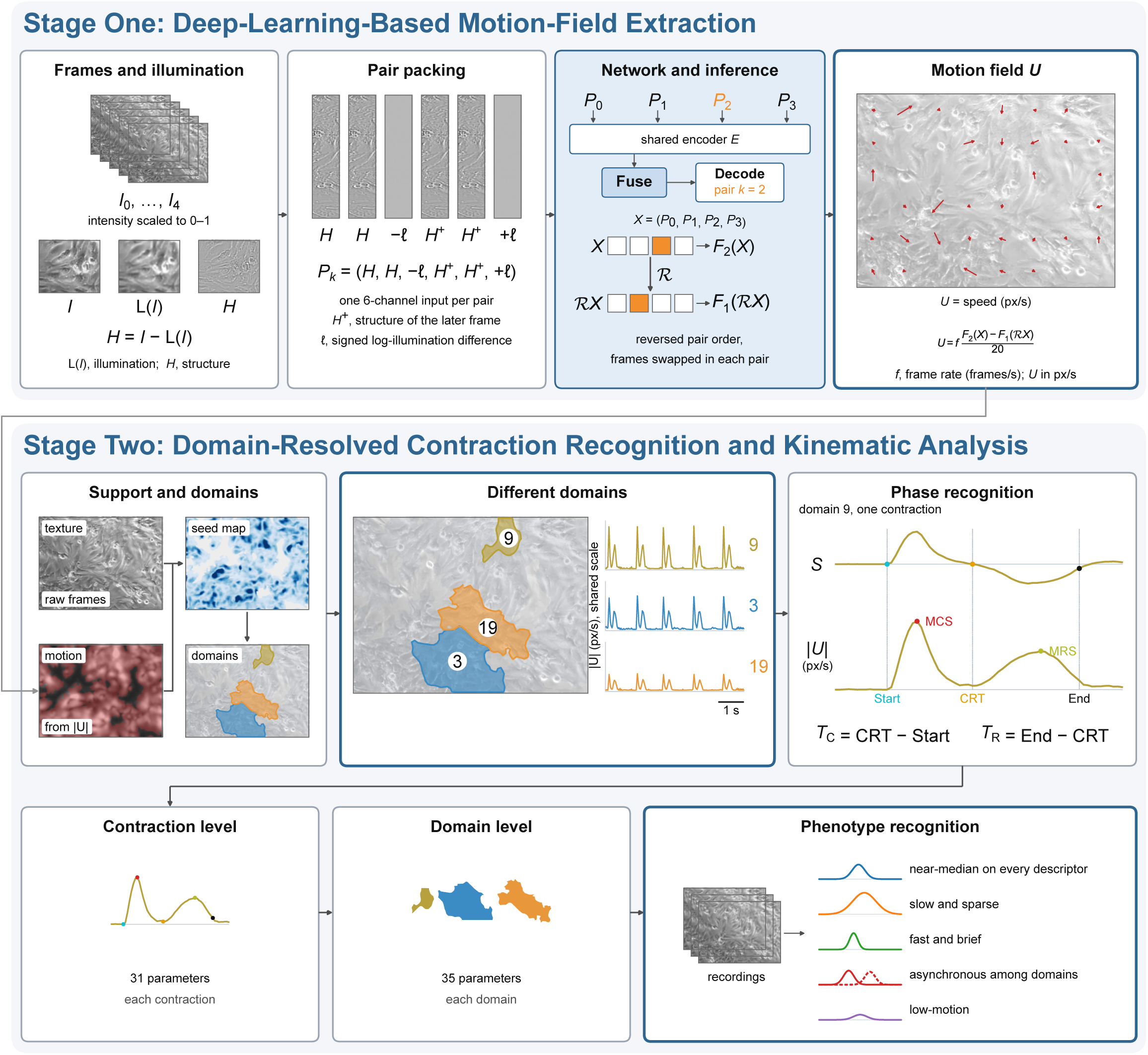
Schematic overview of Cardiomyograph, a label-free method that characterizes contraction from brightfield video. (Top) Stage One estimates the motion field *U* in four steps. (1) Each greyscale frame *I* is split into its blurred copy, the illumination channel, L(*I*), and the structure channel *H* = *I* − L(*I*). (2) Each pair of adjacent frames is packed into a six-channel input *P_k_*. (3) The four pairs of the window *X* are encoded by a shared encoder *E*, fused, and decoded for the orange pair *k* = 2 in a forward pass *F*_2_ (*X*) and a reverse pass *F*_1_ (ℛ*X*). (4) The two passes are combined with the frame rate *f* into *U*, shown on one frame. (Bottom) Stage Two analyses contraction in each domain in four steps. (1) The contractile support, where the texture and motion maps overlap, is partitioned into domains from seeds on the seed map (the mean absolute divergence of *U*). Three of these domains are shown with their |*U*| traces. (2) Contraction and relaxation phases of one contraction of domain 9 are recognized on the traces *S* and |*U*|. (3) Parameters are pooled into contraction-level and domain-level tables. (4) Recording-level features are summarized over the domains of each recording and clustered into contraction phenotypes. Colours and numbers identify the same three domains across boxes, and all images and measured traces are from one recording. Abbreviations: *S* is the directional trace of a domain, positive during contraction and negative during relaxation. |*U*| is the magnitude of *U* averaged over a domain after whole-frame translation is removed. CRT, contraction-relaxation transition; MCS, maximum contraction speed; MRS, maximum relaxation speed; *T*_C_, contraction-phase duration; *T*_R_, relaxation-phase duration.

## Materials and Methods Frame decomposition

### Each greyscale frame *I* is split into a blurred frame *L*(*I*) and a structure channel

*L*(*I*) is a local average of *I* and carries the slowly varying illumination. The structure channel keeps the tissue pattern from which motion is estimated:

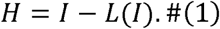

Every frame is scaled to the unit interval before this split. An adjacent pair is then packed as a six-channel input *X*.

### Frame-pair encoding and temporal fusion

Five consecutive greyscale frames form one inference window. The four adjacent pairs in that window are *P_k_*, with pair index *k* = 0,1,2,3. A shared encoder *E* maps the six-channel input of each pair to a feature volume. The four volumes are fused into one representation *Z*, so that the displacement at the selected pair is conditioned on the surrounding frames. Fusion is applied at each spatial scale of the encoder, and the decoder reads the displacement from *Z*. The reported displacement belongs to the third pair, the pair with *k* = 2.

### Forward and reverse inference

The displacement field *U* comes from a forward pass and a reverse pass on the same window. *F*_2_ (*X*) is the prediction for the pair *k* = 2, and *F*_1_ (ℛ*X*) is the prediction for that pair after the window is reversed. The operator ℛ reverses the order of the four pairs and exchanges the earlier frame with the later frame inside each pair. The two predictions are combined antisymmetrically. With *f* the measured frame rate in frames per second, the field in analysis pixels per second is

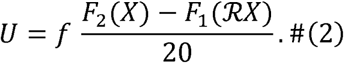

The denominator 20 is the product of the training scale, 10, and the factor 2 from the antisymmetric average. Multiplication by *f* converts the adjacent-frame displacement from analysis pixels into analysis pixels per second. Successive samples of *U* are separated by 1/*f* seconds.

### Synthetic training

Training frames were synthesized from real greyscale textures and from known displacement fields. Each generated frame received a scale, an illumination factor, warping by the known field, and image degradations. The training set contains moving frames and motionless frames.

### Contractile support and domains

The recording is then analysed inside the contractile support Ω, the textured tissue that moves. Background and gaps in the tissue lie outside Ω . Texture is read from a temporal median of greyscale frames, and motion is read from the displacement field *U*. A pixel is admitted to Ω when the texture cue and the motion cue both admit it.

Ω is partitioned into domains Ω_1_, … , Ω*_m_*. A domain is one coherent image region. Markers are placed at local maxima of the positive convergence of *U* at the frames of peak contraction and are grown into domains by a marker-controlled watershed on the negative mean motion magnitude (Supplementary Methods). The partition is computed once for the recording and is then held fixed.

### Contraction-relaxation phase detection

Whole-frame translation is removed before a domain trace is formed. At time *t*, the vector *ξ_t_* is that translation, in analysis pixels per second. For a domain Ω*_j_*, the speed trace |*U*|(*t*) is the mean magnitude of the corrected field inside the domain:

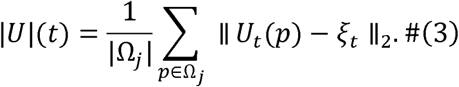

The directional trace *S* follows the dominant motion pattern of the same domain. Translation-corrected vectors inside Ω*_j_* are mean-centred, and *S*(*t*) is the score of their first singular mode:

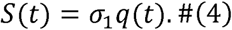

Here *q*(*t*) is the first left singular vector and *σ*_1_ is its singular value. The sign of *S* is set so that contraction is positive and relaxation is negative. One decomposition is held fixed for the whole recording.

Contractions are separated on the displacement obtained by integrating *S* over time. One rise and fall of this displacement is one contraction. Each contraction on these two traces is marked by five landmarks. Start is the time at which *S* rises above a foot level close to its baseline, and End is the time at which *S* comes back within that level of the baseline. The contraction-relaxation transition (CRT) is the time at which *S* crosses zero from positive to negative. The maximum contraction speed (MCS) and the maximum relaxation speed (MRS) are the peaks of |*U*| before and after CRT. The contraction-phase duration *T*_C_ and the relaxation-phase duration *T*_R_ are defined from these landmarks as

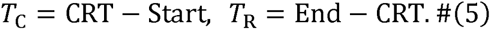

Domains whose contractions coincide are then grouped into synchronous domains, within which contraction rhythm, contraction rate, and synchrony are computed (Supplementary Methods).

### Parameter calculation

Each contraction of each domain supplies one set of 31 contraction-level parameters (Table S1), computed from its landmarks and traces. Each domain supplies one set of 35 domain-level parameters (Table S2). Most domain-level parameters summarize variation among the contractions of that domain.

Further details of the algorithm and Experimental procedures are provided in the Supplementary Material.

## Results

### Ultra-high sensitivity on drug response identification

To assess the performance of Cardiomyograph in detecting drug responses, we treated neonatal rat ventricular myocytes (NRVMs) with a positive inotropic agent, isoprenaline (ISO), at graded concentrations and analysed the resulting videos (Fig. 2). Notably, Cardiomyograph detected a marked increase in contraction amplitude at 10 nM (*P* = 0.014; Fig. 2A-C). The fold change normalized to the dimethyl sulfoxide (DMSO) vehicle rose monotonically from 1.24 at 1 nM to 1.90 at 1,000 nM (Fig. 2D). Additionally, Cardiomyograph readouts agreed closely with Ca^2+^ imaging in contraction amplitude (Pearson *r* = 0.943, *P* < 0.001; Fig. S1B). At 10 nM, Cardiomyograph showed a significant increase in contraction amplitude (Fig. 2D), whereas none of the other three tools showed a significant change (Fig. S2C). Across the dose series, CONTRACTIONWAVE reached significance from 100 nM upwards, while MUSCLEMOTION and Camystat showed no significant response at any tested concentration (Fig. 2E). Within the tested concentration range, the lowest concentration at which Cardiomyograph detected a significant increase in contraction amplitude was at least 10-fold lower than that for the comparison tools (Fig. 2D and 2E).

**Fig. 2.**
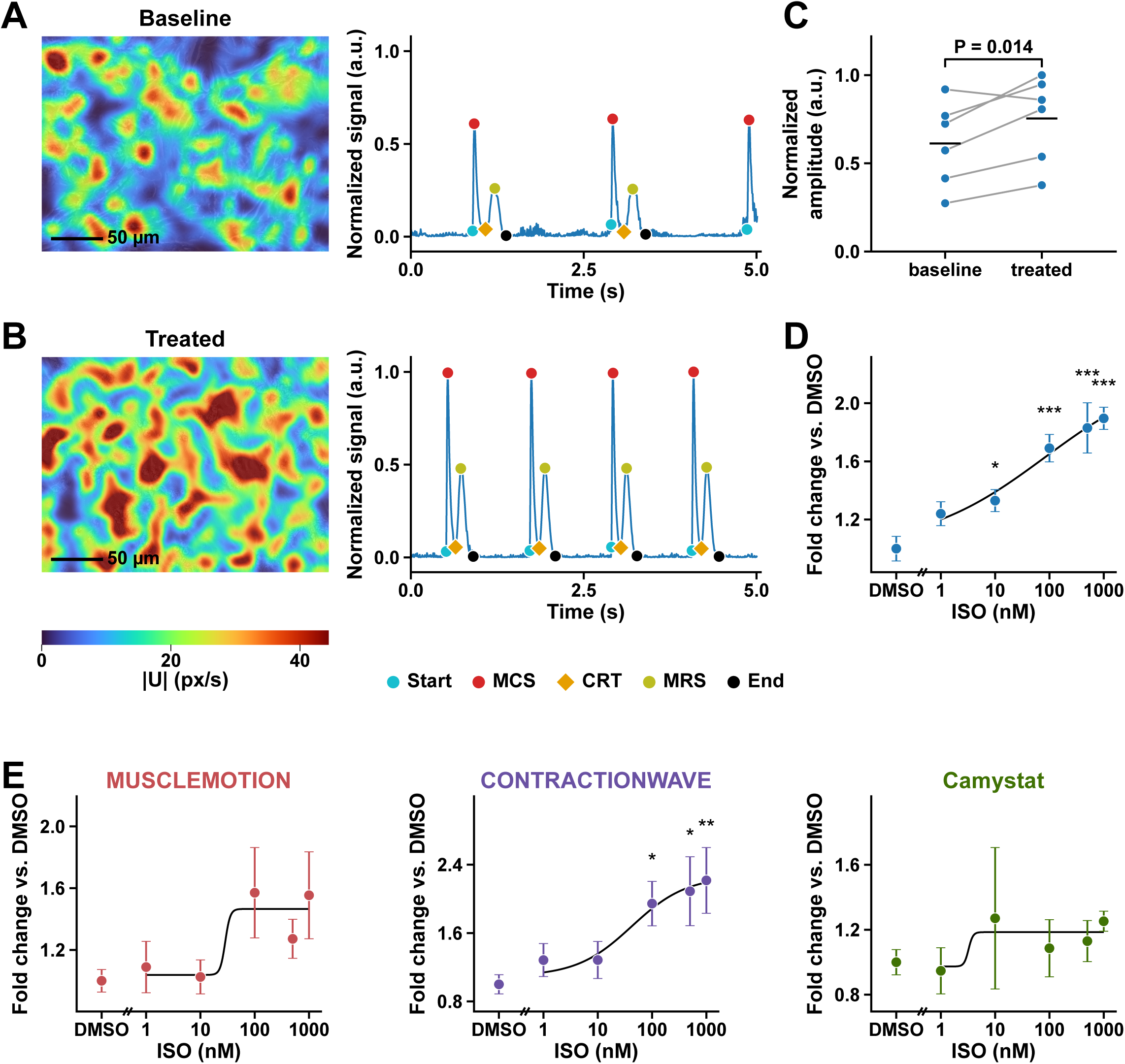
Contraction amplitude in response to isoprenaline, measured by Cardiomyograph and three other label-free contractility tools. (A and B) One well before (A) and after (B) 10 nM isoprenaline (ISO). Left, Cardiomyograph motion-field magnitude |*U*| at the maximum contraction speed (MCS) of one contraction, overlaid on the brightfield frame, with both maps on one colour scale. Right, the whole-field |*U*| signal over 0-5 s windows, min-max normalized by a single factor taken over both windows together. (C) Amplitude of the baseline and treated recordings of the 10 nM wells (*n* = 6), divided by the largest value in the panel. Amplitude is the median, across contractions, of the area of the whole-field |*U*| signal above baseline. Lines join the two recordings of one well, bars indicate group means, and the *P* value is from a two-sided paired t test on log_2_ amplitude. (D and E) Fold change of each well (treated divided by baseline), normalized to the mean of the DMSO wells of the same tool, for Cardiomyograph (D) and for MUSCLEMOTION, CONTRACTIONWAVE, and Camystat (E). Points are mean ± SEM, and curves are four-parameter log-logistic fits. ISO concentration is on a log scale, and DMSO is plotted left of the axis break. Each ISO concentration was compared with the vehicle (0.1% DMSO) wells of the same tool by a two-sided Dunnett test on log_2_ fold change. Scale bars, 50 µm (A and B). *: P < 0.05; **: P < 0.01; ***: P < 0.001. a.u., arbitrary units; CRT, contraction-relaxation transition; DMSO, dimethyl sulfoxide; ISO, isoprenaline; MCS, maximum contraction speed; MRS, maximum relaxation speed.

We next examined the differences among the tools on these recordings. The other three tools showed greater within-recording variability than Cardiomyograph (Fig. S3) and were affected by whole-field drift (Fig. S4), spurious signal at rest and fixed event-detection rules (Figs. S5 and S6). We further benchmarked Cardiomyograph against MUSCLEMOTION, CONTRACTIONWAVE, and Camystat using synthetic cardiomyocyte videos (Fig. S8) and manually annotated bright-field videos with added perturbations (see Supplementary Methods). Spurious motion in static regions was more than 10-fold lower with Cardiomyograph than with published motion-estimation methods (Fig. S7D). Cardiomyograph also achieved higher contraction-detection precision than the other three tools (Fig. S9A). Its mean recall was 0.974, compared with 0.53-0.80 for the other three tools (Fig. S9B). Across contraction onset, the contraction-relaxation transition, and the end of contraction, the median absolute timing error on the annotated recordings was 5.7 ms for Cardiomyograph and 41.7-120.7 ms for the other three tools (Fig. S9C-E). These findings indicate that Cardiomyograph was less susceptible to background signal and drift.

A similar advantage in dose-response sensitivity was observed with two negative inotropic agents, lacidipine and verapamil. These results show that Cardiomyograph can sensitively and accurately quantify drug-induced changes in cardiomyocyte contraction.

### Precise quantification of the diastolic phase in cardiomyocyte contractions

Contraction and relaxation durations are key temporal parameters for assessing cardiomyocyte mechanical function, and precise determination of the onset and end of each contraction and relaxation is particularly critical. However, existing video-based tools remain limited in reliably delineating relaxation. To determine whether Cardiomyograph could accurately resolve the onset and end of contraction and relaxation, especially the relaxation phase, we established a cell model of prolonged relaxation using E-4031 (50 nM), which delays repolarization by blocking the rapid component of the delayed rectifier K^+^ current, IKr^17^. We then compared Cardiomyograph measurements of relaxation duration with those obtained using three existing tools. Both Cardiomyograph and MUSCLEMOTION could detect E-4031-induced prolongation of relaxation, whereas CONTRACTIONWAVE did not detect a significant change and the Camystat comparison, based on only two paired recordings, was inconclusive (Fig. 3). These two tools failed mainly because they could not accurately locate the end of each contraction (Fig. S10D and S10E). Although MUSCLEMOTION responded sensitively to the change in relaxation duration, its measurements showed greater within-recording dispersion and lower stability than those of Cardiomyograph. The median within-recording coefficient of variation was 0.25 for MUSCLEMOTION and 0.045 for Cardiomyograph (Fig. S10C).

**Fig. 3.**
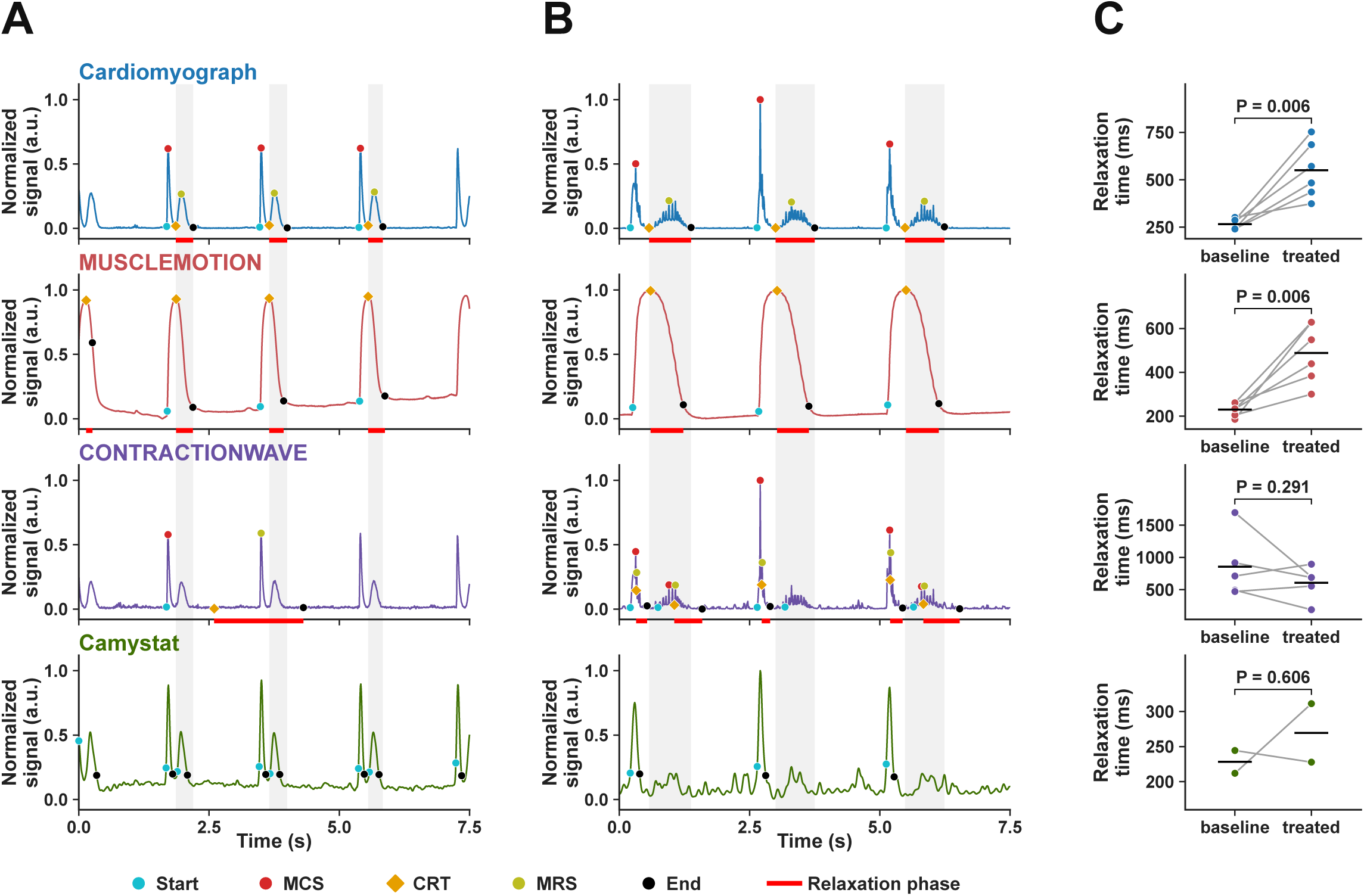
Detection of functional changes in the contractile time course. (A and B) Contraction traces before (A) and after (B) E-4031 (50 nM) from Cardiomyograph, MUSCLEMOTION, CONTRACTIONWAVE, and Camystat (top to bottom). Markers are landmarks from each tool’s own output. Grey bands span the contraction-relaxation transition (CRT) to End of each marked Cardiomyograph contraction and extend across all four tools. Red bars below the Cardiomyograph, MUSCLEMOTION, and CONTRACTIONWAVE traces mark the relaxation phases detected by each tool. (C) Paired relaxation times for each tool (rows as in A and B). Each dot is the median relaxation time of one recording. Lines connect paired recordings, and horizontal bars indicate group means. *P* values are from a two-sided paired t test. a.u., arbitrary units; CRT, contraction-relaxation transition; MCS, maximum contraction speed; MRS, maximum relaxation speed.

Furthermore, we investigated whether Cardiomyograph could resolve the temporal course of contraction and relaxation in models of irregular contraction rhythm induced by oxygen-glucose deprivation (OGD) and doxorubicin (DOX). Cardiomyograph captured the timing of every irregular contraction event without missing a single contraction (Fig. S11A and S11B). In the DOX experiment, Cardiomyograph detected the same number of contractions as manual review in 98% of recordings, compared with 33%-83% for the other three tools (Fig. S11C). The rhythm grade derived from its contractions matched the manual grade in 98% of recordings, compared with 51%-93% for the other three tools (Fig. S11D). Together, these findings indicate that Cardiomyograph effectively delineates the temporal boundaries of contraction and relaxation phases, enabling reliable extraction of temporal features.

### Uniquely identification of ectopic pacemakers and analysis of contraction heterogeneity

Cultured cardiomyocytes frequently develop ectopic pacemaking during differentiation and maturation. Across videos of NRVMs and human induced pluripotent stem cell-derived cardiomyocytes (hiPSC-CMs), focal contractions were detected in 826 of 2,647 videos with contractions (31.2%; Fig. S12). However, since MUSCLEMOTION, CONTRACTIONWAVE and Camystat generate whole-field readouts by averaging motion across the entire field of view, regional contractile heterogeneity was obscured by these tools. Their readouts can neither localize ectopic sources nor provide region-specific measurements (Fig. 4A).

**Fig. 4.**
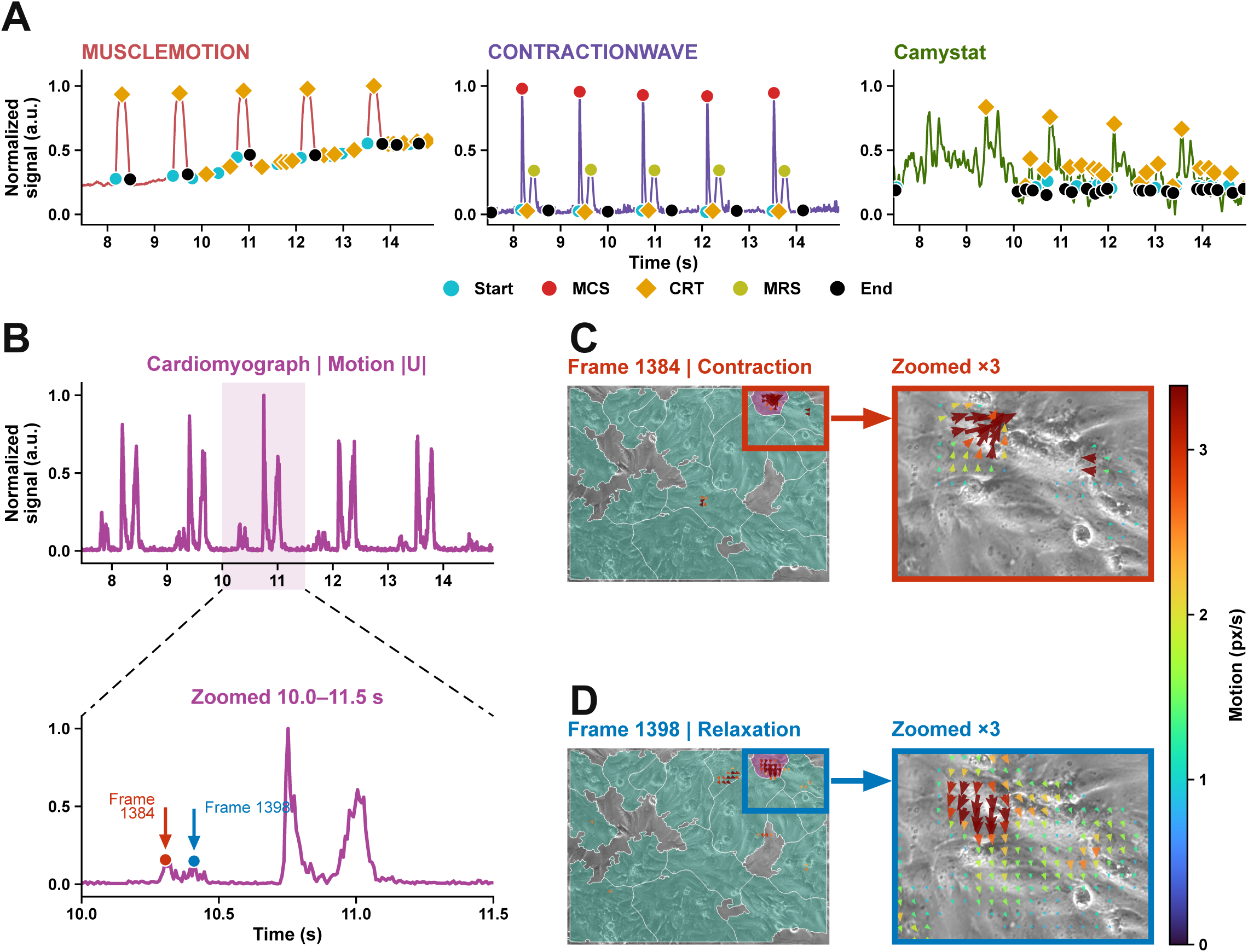
Detection of a focal abnormal contraction via domain-level motion mapping. (A) MUSCLEMOTION, CONTRACTIONWAVE, and Camystat output. Each trace is normalized to 0-1. Markers are landmarks from each tool’s own output. For MUSCLEMOTION and Camystat, they show Start, End, and the signal peak, labelled CRT; for CONTRACTIONWAVE, they also show MCS and MRS, and CRT marks the speed minimum between them. Camystat parameters were adjusted on this recording, with the MUSCLEMOTION peaks as the reference. (B) Cardiomyograph output. The motion signal |*U*| of the focal domain is normalized to 0-1 and shown for the second half of the recording (top) and enlarged over 10.0-11.5 s, the window marked by the shaded band (bottom). The red arrow marks frame 1384, during the focal contraction, and the blue arrow marks frame 1398, during the following relaxation. (C and D) Displacement fields at frames 1384 (C) and 1398 (D). On the full field, the domain that produces the focal contractions is shown in purple and the other domains in teal. The boxed region is enlarged 3×. Arrows give the local direction of motion, and colour gives the magnitude in px/s (scale at right). a.u., arbitrary units; CRT, contraction-relaxation transition; MCS, maximum contraction speed; MRS, maximum relaxation speed.

Cardiomyograph addresses this limitation by automatically identifying contractile domains and extracting motion parameters independently for each domain (Fig. S13). In a representative recording, Cardiomyograph partitioned the field into 17 domains and identified a focal region occupying 2.0% of the analysed area. The trace revealed a focal contraction before the next field-wide contraction (Fig. 4B). Displacement fields at frames 1384 and 1398 captured contraction and relaxation of the focal region, respectively (Fig. 4C and 4D). Two additional videos showed the same pattern, with a focal domain among 23 and 17 domains, respectively (Fig. S14). Among the evaluated methods, Cardiomyograph was the only approach that localized focal regions masked by field-wide averaging and quantified their motion on a domain-by-domain basis.

Cardiomyograph therefore provides a spatially resolved approach to quality control of cardiomyocyte preparations. This capability could improve the reliability of downstream drug screening and support mechanistic studies of diseases associated with asynchronous contraction.

### Specific diagnosis of abnormal waveforms

To evaluate the applicability of Cardiomyograph across diverse experimental settings, we analysed videos of hiPSC-CMs and a cardiac organoid, as well as videos acquired using differential interference contrast microscopy and traction force microscopy (Fig. 5A). Across these preparations and imaging modalities, Cardiomyograph consistently generated waveforms that clearly delineated contraction dynamics.

**Fig. 5.**
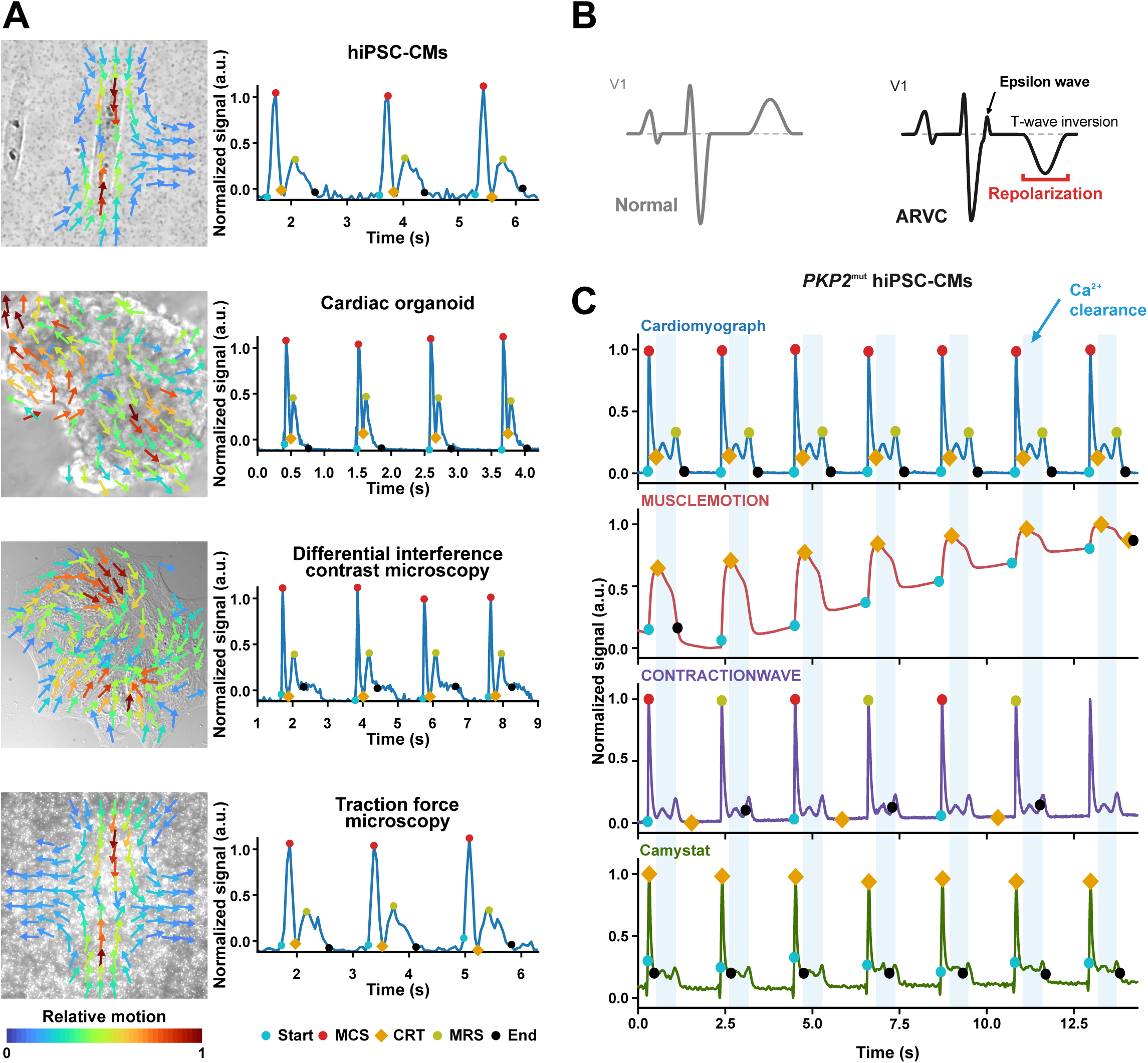
Application of Cardiomyograph across multiple cardiac preparations and imaging modalities. (A) Representative motion fields and corresponding normalized Cardiomyograph contraction-speed traces from, top to bottom, human induced pluripotent stem cell-derived cardiomyocytes (hiPSC-CMs), a cardiac organoid, differential interference contrast microscopy, and traction force microscopy. Arrows give the local direction of motion, and colour gives its relative magnitude. Symbols are shown only for complete contractions. (B) Schematic lead V1 electrocardiograms under normal conditions and in arrhythmogenic right ventricular cardiomyopathy (ARVC). (C) Contraction traces from one recording of hiPSC-CMs carrying a heterozygous *PKP2* variant (ARVC hiPSC-CMs), obtained with Cardiomyograph, MUSCLEMOTION, CONTRACTIONWAVE, and Camystat, each min-max normalized over the recording. Markers are landmarks from each tool’s own output. Blue bands, labelled Ca^2+^ clearance, span CRT to MRS of each Cardiomyograph contraction and extend across all four tools. a.u., arbitrary units; ARVC, arrhythmogenic right ventricular cardiomyopathy; CRT, contraction-relaxation transition; hiPSC-CMs, human induced pluripotent stem cell-derived cardiomyocytes; MCS, maximum contraction speed; MRS, maximum relaxation speed; PKP2, plakophilin-2.

Cardiomyopathies caused by pathogenic genetic variants can manifest clinically as abnormalities in cardiac contraction. We therefore investigated whether such variants could also give rise to detectable abnormalities in cultured cardiomyocytes. To this end, we recorded videos of arrhythmogenic right ventricular cardiomyopathy (ARVC) hiPSC-CMs^18^. Characteristic electrocardiographic findings in ARVC include T-wave inversion (a repolarization abnormality) and epsilon waves (a depolarization abnormality) in the right precordial lead (Fig. 5B)^19^. Cardiomyograph resolved the abnormal contractile dynamics in ARVC hiPSC-CMs, a slower transition from contraction to relaxation (Fig. 5C), consistent with the prolonged Ca^2+^ transients reported in plakophilin-2 (PKP2)-deficient models^20, 21^. In contrast, due to the complexity of cellular contractile activity, none of the three tools segmented the contraction cycle correctly or completely, or delineated the phase from the contraction-relaxation transition (CRT) to the maximum relaxation speed (MRS) in individual contractions, though MUSCLEMOTION detected some fluctuations (Fig. 5C). This result can be attributed to the fixed rules that all three tools apply to a single averaged trace, such as peak order or intensity thresholds, which do not hold for complex waveforms.

Together, these results demonstrate the advantage of Cardiomyograph in analysing complex contractile waveforms and suggest its potential for phenotypic analysis and prediction using *in vitro* models of genetic cardiomyopathy.

## Discussion

Cardiomyocyte contraction is essential for the pumping function of the heart which drives blood circulation, maintains organ perfusion, and determines cardiac output. Here, we introduce Cardiomyograph, a label-free deep-learning framework that addresses two barriers in video-based contractile analysis, unreliable motion representation and the loss of spatial and directional information during signal aggregation. Our results demonstrate that Cardiomyograph effectively quantifies cardiomyocyte contractile dynamics and generates interpretable functional reports that can be used for routine assessment and screening. Cardiomyograph generates stable parameters without manual tuning, reliably identifies asynchronously contractile regions and sensitively detects subtle pharmacological perturbations.

Conventional approaches, including patch clamp, calcium imaging, microelectrode arrays and traction force microscopy, provide valuable mechanistic insights into cardiomyocyte function but their broader application is constrained by invasiveness, experimental complexity, and high operational cost. Video-based approaches have therefore emerged as attractive alternatives because they enable noninvasive analysis of microscopy recordings^10, 11, 22^. However, most existing methods remain highly sensitive to noise, baseline drift, and platform heterogeneity due to the dependence on one-dimensional signals and manual parameter tuning. Although field-wide signals can summarize overall activity, contraction-relaxation events from different regions are merged and weak domains are obscured. There are two major limitations in current video-based contractility analysis. One concern is inaccuracy in motion representation.

Frame-difference methods infer activity from changes in image intensity^10–12^, optical-flow methods estimate pixel displacement^13, 14^, and image-correlation methods match small image windows between frames^15, 16^. All three can be affected by noise, drift of the cell layer, floating debris and illumination changes. The other limitation is that existing video-based analysis discards spatial motion information when local measurements are aggregated into a single activity trace. As a result, the directional information of the motion vectors is lost, which in turn makes the identification of the contraction process and its phases inaccurate and obscures regional differences. Thus, phase assignment remains sensitive to decision rules and user-selected thresholds.

These limitations entail two direct consequences for disease modelling and drug evaluation. First, inaccurate detection of contraction-relaxation phases leads to temporal mislocalization of contraction events. As a result, estimates of contraction rate and the detection of missed contractions become unreliable, while amplitude, velocity, and duration are computed over incorrectly defined temporal windows^10, 23^. Effects confined to a single phase, such as impaired diastolic relaxation in hypertrophic cardiomyopathy^24^ or prolonged systole mediated by cardiac myosin activators^25^, may likewise remain undetected. Second, whole-field averaging obscures regional heterogeneity in contractile behaviour across the culture. For example, strain values can vary substantially even between adjacent regions of a cardiomyocyte monolayer^26^. Electrical coupling between adjacent cells may be spatially heterogeneous or attenuated, thereby reducing impulse conduction velocity^4^ and increasing susceptibility to arrhythmogenesis^27^. Following focal injury to engineered cardiac microtissues, global contractile force recovers to baseline levels owing to compensation by uninjured regions, whereas the injured region exhibits complete contractile failure that is undetectable by whole-tissue force assays^28^.

Cardiomyograph addresses these issues through improved motion-field estimation, regional analysis, and complementary directional information. At the image level, temporal context, a separate illumination channel, and zero-motion supervision help distinguish coherent deformation from artefacts. At the beat level, the directionality of the motion vectors assists the identification of the contraction process and its phases, while motion magnitude characterizes activity strength.

Cardiomyograph enables stable generalization across diverse datasets without manual adjustment. Existing tools either tune parameters for each recording, which introduces operator bias, or apply a single parameter set to a whole batch at the cost of accuracy on heterogeneous recordings. Unlike representative existing tools such as MUSCLEMOTION, CONTRACTIONWAVE and Camystat, which are vulnerable to static-background interference, including illumination changes, contractile-signal dilution and false motion detection in background regions, Cardiomyograph employs adaptive contractile-domain detection to autonomously isolate contractile regions and thereby derives motion traces that more faithfully represent true contractile activity. This improves parameter stability and helps overcome the reproducibility limitations of pixel-intensity or optical-flow-based methods. Cardiomyograph is also capable of localizing regions that fail to synchronize during maturation, enabling further study of their features. The drug-sensitivity test highlights the sensitivity of Cardiomyograph to subtle pharmacological perturbations, and the ARVC experiment highlights its accuracy in resolving complex contractile waveforms. In ARVC hiPSC-CMs, Cardiomyograph resolved the slowed contraction-to-relaxation transition, whereas the other three tools did not delineate this phase.

Nevertheless, several limitations of the present work should be acknowledged. First, Cardiomyograph relies on two-dimensional video recordings and therefore cannot directly resolve through-plane motion. In future work, three-dimensional light-sheet volumetric imaging will be used to recover the through-plane motion components that are collapsed in two-dimensional projections. Second, although Cardiomyograph generates interpretable phenotype reports, disease-specific classification has not yet been validated in independent clinical cohorts. Our next step will be to establish multicenter, independently labelled cohorts of patient-derived hiPSC-CMs.

Electrocardiography (ECG) is an established tool for assessing cardiac disease. EchoNet-Dynamic^29^ and EchoNext^30^ demonstrate how echocardiographic videos and ECG recordings, respectively, can be linked to defined functional and disease-related targets. However, translating contraction-relaxation dynamics into well-defined disease-associated phenotypes at the cellular level remains challenging. Cardiomyograph supports continuous characterization of local mechanical behaviour and is capable of relating contractile activity to disease-associated phenotypes, while the use for disease-specific classification requires further development and validation. Future studies should extend these measurements by developing a dedicated classifier on large cohorts with independently established disease labels and evaluating its performance in independent datasets, which may relate contractile phenotypes to ECG and echocardiographic findings. Cardiomyograph will accelerate mechanism dissection and drug development through enabling high-throughput screening of specific contractile phenotypes that are difficult to capture with conventional global traces.

## Conclusions

In summary, Cardiomyograph is a label-free deep-learning platform that effectively quantifies cardiomyocyte contractile dynamics and offers interpretable functional phenotype reports, supporting routine culture assessment, longitudinal monitoring and rapid large-scale drug screening.

## Sources of Funding

This study was supported by the National Natural Science Foundation of China (grant Nos. 82630122, 82425060 and 82270350 to Y. Xiang, 82670508 and 82300326 to Y. Xu, and 82300381 to K. Jiang) and the Noncommunicable Chronic Diseases-National Science and Technology Major Project (grant Nos. 2025ZD0546900 and 2025ZD0546901 to Y. Xiang).

## Data, Materials and Software Availability

Detailed information on materials, methods and software implementation is available at https://github.com/XianglabHeart/Cardiomyograph.

## Author contributions

Y. Xiang designed the study. X. Yuan, K. Li and Q. An performed the experiments. Y. Xu, X. Yuan and D. Wang analysed the data. X. Yuan, K. Li, Y. Xu, D. Wang, K. Jiang, J. Wang and Y. Xiang wrote the paper. All authors read and approved the final manuscript.

## Disclosures

The authors declare no competing interests.

## Abbreviations

ARVC: arrhythmogenic right ventricular cardiomyopathy
CRT: contraction-relaxation transition
DMSO: dimethyl sulfoxide
DOX: doxorubicin
hiPSC-CMs:: human induced pluripotent stem cell-derived cardiomyocytes
ISO: isoprenaline
MCS: maximum contraction speed
MRS: maximum relaxation speed
NRVMs: neonatal rat ventricular myocytes
OGD: oxygen-glucose deprivation

## Notes

### Competing Interest Statement

The authors have declared no competing interest.

