## Supplementary Materials for "Cardiomyograph: a robust tool for evaluating contraction of cardiomyocytes and cardiac organoids with ECG-like reports"

**This file includes:**

Supplementary Methods

Figures S1 to S14

Tables S1 and S2

### Supplementary Methods

#### Algorithm details

##### Frame decomposition

Each adjacent pair is packed into $X$ as follows. The structure channel of the earlier frame is entered twice, the structure channel $H^{+}$ of the later frame is entered twice, and the illumination difference $\mathcal{l}$ between the two frames is entered once with each sign:

$$\begin{aligned} X=(H,H,-\mathcal{l},H^{+},H^{+},+\mathcal{l}).\#\left( 6 \right) \end{aligned}$$

Here $\mathcal{l}$ is the signed log-illumination difference of the pair. The repeated structure channels give the shared encoder a cue for tissue motion, and the two signed illumination channels give it a cue for lighting. Every adjacent pair used to estimate the displacement field is packed in this way.

##### Frame-pair encoding and temporal fusion

For a window centred on frame $I_{i}$, the pair $k=2$ is the step from $I_{i}$ to the next frame. The other three pairs are encoded with the same encoder and enter only through the fused representation $Z$. The field returned for the window is the field of this third pair.

##### Synthetic training

With probability 0.5, each training window, motionless or moving, also received a whole-frame translation of 0.03-0.6 analysis pixels per frame. On a motionless frame without this translation, the target displacement is zero. The known field was scaled and then accumulated across the five frames of the window, and the pair $k=2$ carried the displacement that inference reads out.

The loss has three terms. $J_{\mathrm{pair}}$ penalizes the displacement error of each pair, $J_{\mathrm{sum}}$ penalizes the error of the four pair outputs added together, and $J_{\log}$ penalizes the error of log amplitude:

$$\begin{aligned} J=J_{\mathrm{pair}}+2J_{\log}+2J_{\mathrm{sum}}.\#\left( 7 \right) \end{aligned}$$

##### Contractile support and domains

The outer 8 analysis pixels of the image content are excluded from the support. The seed map is the positive convergence of $U$ (the negative divergence, with negative values set to zero), computed after a Gaussian of 10 analysis pixels is applied to the field and again to the divergence. It is averaged over the frames of peak contraction, each weighted by the mean convergence on the support in that frame. A frame of peak contraction is the frame with the highest mean convergence in each run of frames in which this mean reaches half of its maximum over the recording. Markers are placed at local maxima of the seed map, in a 25-pixel window and at least 15 pixels apart, on support pixels that lie at least 32 analysis pixels from the edge of the image content. Markers are merged when their basins meet above the 75th percentile of the positive values of the seed map on those pixels. The mean motion magnitude is taken over the frames that remain once the 1% of frames with the highest whole-frame motion are set aside and the next 10% are kept. The merged markers grow by a marker-controlled watershed on the negative of the mean motion magnitude of those same frames, smoothed by a Gaussian of 5 analysis pixels. Domains smaller than one marker disc, $\pi\times{15}^{2}\approx707$ analysis pixels, are removed, and their pixels remain in the support without a domain label.

##### Synchronous domains

Two domains are grouped into one synchronous domain when at least 90% of their main contractions coincide within 60 ms. A main contraction reaches at least 0.3 of the typical swing of $S$ and of the typical speed of its domain. The inter-contraction intervals in Table S2 are taken in the synchronous domain to which a domain belongs, and contractions truncated by the start or end of the recording are excluded.

##### Parameter calculation

All five landmarks are located between frames. Start and End are linearly interpolated where $S$ crosses the foot level, and CRT where $S$ crosses zero. The foot level is the larger of twice the noise SD of $S$ and 2% of the peak of $S$ in that contraction, and is at most the half-width of the baseline band. MCS is the earliest local peak of $|U|$ between Start and CRT that reaches 80% of the highest such peak, and MRS is the highest value of $|U|$ between CRT and End. Both are placed at the vertex of a parabola through the peak sample and its two neighbours. In Tables S1 and S2, $\tau_{0}$, $\tau_{2}$, and $\tau_{4}$ are the fractional frame positions of Start, CRT, and End, and $t_{0}$, $t_{2}$, and $t_{4}$ denote the same positions.

Expressed in milliseconds, the durations $T_{C}$ and $T_{R}$ of Eq. 5 and the onset-to-end duration $T_{B}$ (Table S1, rows 16, 17, and 1) are

$$\begin{aligned} T_{C}=\frac{1000(\tau_{2}-\tau_{0})}{f}, T_{R}=\frac{1000(\tau_{4}-\tau_{2})}{f}, T_{B}=\frac{1000(\tau_{4}-\tau_{0})}{f}.\#\left( 8 \right) \end{aligned}$$

The contraction speed integral $\Lambda$ (Table S1, row 24), in analysis pixels, is the part of $|U|$ that lies above the lower of its values at Start and CRT, integrated over time from Start to CRT by the trapezoidal rule. With $t$ in frames,

$$\begin{aligned} \Lambda=\frac{1}{f}\int_{\tau_{0}}^{\tau_{2}} \max(|U|(t)-min(|U|(\tau_{0}),|U|(\tau_{2})),0) dt.\#\left( 9 \right) \end{aligned}$$

The contraction rate, in $\min^{-1}$, is 60 divided by the median inter-contraction interval in seconds (Table S2, row 26). Speed at MCS is $|U|(t_{\mathrm{MCS}})$ (Table S1, row 22). Displacement is the time integral, from Start, of the mode score before its temporal mean is removed. The time to peak displacement (Table S1, row 3) is the time from Start to the peak of this displacement within the contraction. The 50% displacement recovery time (Table S1, row 2) is the time from that peak until the displacement first falls halfway from the peak to its value at End, with the crossing interpolated between frames.

Missing values are excluded from any summary that requires them. The duration of a complete contraction uses all five landmarks in temporal order. When the relaxation phase is truncated, contraction-phase quantities (e.g., $T_{C}$ and MCS) remain available, whereas the relaxation-dependent quantities are excluded. A summary of the relative interquartile range (rIQR) uses at least five finite contractions. A summary of the coefficient of variation (CV) uses at least eight finite contractions. A rhythm summary uses at least three complete contractions.

#### Benchmarks

Synthetic recordings were generated by warping a greyscale texture from a real recording with a known displacement field, so the frame-to-frame ground truth is known. Twelve perturbations were each applied at four levels to five textures: reduced contraction amplitude, noise, dimming, defocus, slow drift, shaking, a change of frame rate (to 134, 90, 60, or 30 frames per second), combined dimming and noise (with slow drift from the second level and brightness flicker at the fourth), video compression, uneven illumination, debris in the field, and saturation. Together with one unperturbed recording per texture, this gave 245 recordings (Figs. S7 and S8). For the real benchmark, reference labels were produced by manual editing of the events detected by Cardiomyograph, with its motion field and curves visible. Ten of the 40 annotated recordings were drawn at random, and 12 perturbed versions of each were generated, giving 120 recordings (Fig. S9). The perturbations were added Gaussian noise, brightness and contrast scaled to 0.80, Gaussian blur (σ of 2.0 or 3.5 camera pixels), sinusoidal drift (peak shift of 0.4 or 1.0 camera pixels within 45 ms), shaking at 7.3 Hz, resampling to 60 frames per second, uneven illumination (amplitude of 0.15 or 0.30), and saturation of 2% or 8% of the pixels.

##### Motion fields

The released Cardiomyograph field was compared with DEFORM-Net^1^, CONTRACTIONWAVE^2^ using Farnebäck optical flow^3^, and OpenCV dense inverse search (DIS)^4^. All four methods received the same analysis images. Let $\hat{u}(p)$ be the estimated adjacent-frame displacement and $u(p)$ the reference displacement, both in analysis pixels at pixel $p$. Endpoint error (EPE) and direction error are

$$\begin{aligned} EPE(p)=\parallel\hat{u}(p)-u(p)\parallel_{2}, \vartheta(p)=arccos\left( \frac{\hat{u}(p)\cdot u(p)}{\parallel\hat{u}(p)\parallel\parallel u(p)\parallel} \right).\#\left( 10 \right) \end{aligned}$$

The direction error is expressed in degrees and is evaluated only where the reference displacement is at least 0.0125 analysis pixels. Spurious motion was measured in the 190 recordings that contain a static region.

##### Comparison tools

MUSCLEMOTION v1-1beta was used from https://github.com/l-sala/MUSCLEMOTION^5^. CONTRACTIONWAVE was used from https://github.com/marceloqla/ContractionWavePy^2^. Camystat comparisons used a port whose acquisition stage follows Camystat v1.0.4 and whose post-processing follows v1.0.5-alpha-2 (https://github.com/camystat/camystat)^6^. Each tool was run with its native readout: MUSCLEMOTION measures the intensity difference between each frame and a reference frame, CONTRACTIONWAVE computes Farnebäck optical flow, and Camystat counts pixels whose binarized value changes between consecutive frames. For each tool, amplitude was the median, across contractions, of the contraction amplitude reported by MUSCLEMOTION, of the area of the CONTRACTIONWAVE speed curve above baseline, and of the raw Camystat percentage peak within each event window. For each comparison tool, a set of candidate settings that included the default was fixed before scoring, and the setting with the highest F1 score was used. For Fig. S9, it was selected on the 40 annotated recordings without perturbation. In the ISO and DOX experiments, it was selected against manually reviewed contractions. Cardiomyograph used one fixed set of settings for all recordings and was never tuned per recording.

##### Contraction events

A detected event matches a reference event when its representative time lies within 0.35 s of the reference CRT. The representative time is the CRT for Cardiomyograph, the peak for MUSCLEMOTION, the speed minimum between the two speed maxima for CONTRACTIONWAVE, and the maximum of the smoothed trace for Camystat. Matches are assigned one-to-one. Precision, recall, and their harmonic mean, the F1 score, are

$$\begin{aligned} Precision=\frac{N_{\mathrm{match}}}{N_{\mathrm{tool}}}, Recall=\frac{N_{\mathrm{match}}}{N_{\mathrm{ref}}}, F1=\frac{2N_{\mathrm{match}}}{N_{\mathrm{tool}}+N_{\mathrm{ref}}}.\#\left( 11 \right) \end{aligned}$$

$N_{\mathrm{match}}$, $N_{\mathrm{tool}}$, and $N_{\mathrm{ref}}$ count matches, tool events, and reference events. Timing errors use the time points reported by each tool itself.

#### Public microscopy recordings

The single-cell bright-field and traction-force-microscopy (TFM) examples of human induced pluripotent stem cell-derived cardiomyocytes (hiPSC-CMs) were obtained from a published study^7^. The organoid example was obtained from the supplementary material of MUSCLEMOTION^5^. The differential-interference-contrast (DIC) example came from the cell1_dicm image sequence of DEFORM-Net^1^. These examples are published recordings reused for analysis. The DIC source consists of 750 TIFF images. Analysis preserves the supplied frame order and the time base of the actual input movie. Motion inferred from the TFM movie is reported as image displacement.

#### Neonatal rat ventricular myocyte isolation and culture

Neonatal rat ventricular myocytes (NRVMs) were isolated from Sprague-Dawley (SD) neonatal rats. Neonatal rats were deeply anaesthetized with isoflurane and disinfected with 75% ethanol. Hearts were excised, and ventricular tissue was minced and digested overnight at 4 °C in 0.25% EDTA-free trypsin (Gibco, #15050065). After the trypsin was removed, the tissue was digested with 0.1% type II collagenase (Gibco, #17101-015) at 37 °C for 15 min, and this step was repeated four times until digestion was complete. The combined supernatant was filtered through a 100-µm mesh and centrifuged at 800 rpm for 5 min. The cell pellet was resuspended in 1× red blood cell lysis buffer (BioLegend, #420302), held for 5 min, and centrifuged again. Cells were cultured in F12/DMEM (1:1) medium (Gibco, #11330032) supplemented with 10% serum (Gibco, #A5669701) and 1% penicillin-streptomycin (Gibco, #15140122) at 37 °C under 5% CO₂ for 1.5 h to allow differential attachment. The supernatant, which contained the cardiomyocytes, was collected, and cells were plated in 24-well plates at 0.8 × 10^6^ cells/mL. After 36-48 h, once spontaneous beating was observed, the medium was replaced with F12/DMEM (1:1) containing 10% serum, 1% penicillin-streptomycin, and 100 µM BrdU (Sangon, #A417150-0500) to suppress the proliferation of residual fibroblasts, and the medium was then changed every 2 days.

#### hiPSC-derived cardiomyocytes

HiPSCs (CTCC-001-0783, Zhejiang Meisen Cell Technology Co., Ltd.) were differentiated into cardiomyocytes by a published protocol^8^ with minor modifications: 5 µM CHIR99021 (STEMCELL, #77054) was added on day 0, and 5 µM IWP2 (Tocris, #3533) and 5 µM XAV939 (Sigma, #X3004-5MG) were added on day 3. Thereafter, RPMI 1640 GlutaMAX medium (Gibco, #61870036) containing B27 minus insulin (Gibco, #A1895601) was changed every 2 days.

Commercially differentiated, supplier-designated arrhythmogenic right ventricular cardiomyopathy (ARVC) hiPSC-CMs purchased from Nanjing Cosmos Biotechnology Co., Ltd. were maintained separately. Cells were plated at 1×10^6^ cells per well in 24-well plates. The accompanying recovery, plating, and maintenance media were CSM-H230012, CSM-H230014, and CSM-H230015, respectively. After thawing and recovery, the cells were seeded in plating medium. Two days later, the medium was replaced by maintenance medium, which was subsequently changed every two days.

#### Experimental conditions and perturbations

##### Baseline

Before imaging, the medium was exchanged to remove floating cells, which would otherwise contaminate the contraction analysis. Beating was recorded 1-4 h after the medium change to establish the baseline, and treatments were applied afterwards.

##### Positive inotropic agent

Isoprenaline (ISO; MCE, #HY-B0468), a positive inotropic agent, was diluted in culture medium to 1, 10, 100, 500, and 1,000 nM. The final dimethyl sulfoxide (DMSO) concentration was adjusted to 0.1% (v/v) in all wells. Cells were washed twice with pre-warmed phosphate-buffered saline (PBS), the ISO-containing medium was added, and the cells were cultured at 37 °C for 1 h before recording.

##### Repolarization-delaying agent

E-4031 (MCE, #HY-15551), an agent that delays repolarization, was applied to NRVMs at 50 nM. The final DMSO concentration was adjusted to 0.1% (v/v) in all wells. Cells were washed twice with pre-warmed PBS, the E-4031-containing medium was added, and the cells were cultured at 37 °C for 15 min before recording.

##### Oxygen-glucose deprivation

Cells were washed twice with PBS pre-warmed to 37 °C. Glucose-free DMEM (Gibco, 11966025), containing 2-3% serum, was added. Cells then underwent oxygen deprivation at 37 °C for 20 min before recording.

##### Cardiotoxic agent

Doxorubicin (DOX; MCE, #HY-15142) was applied to NRVMs at 0.1, 0.3, 0.5, 1, and 2 µM. Control wells received the solvent alone. Each field of view was recorded before treatment and after 20 h of exposure. The contractions detected in the DOX recordings were reviewed manually to give the reference number of contractions and rhythm grade. Rhythm was graded as irregular when the coefficient of variation of the inter-contraction intervals was at least 0.25, as regular when it was below 0.25, and as too few contractions when fewer than three were detected.

#### In-house microscopy recordings

Bright-field recordings were acquired on an inverted microscope (Olympus) with a 20× objective and a high-speed digital CMOS camera (Point Grey Grasshopper3, GS3-U3-51S5M-C) controlled by FlyCapture2 Camera Selection 2.13.3.61. One camera pixel corresponds to 0.1725 µm. Frames of 1,600 × 1,200 pixels were downsampled twofold to 800 × 600 pixels for analysis, so that one analysis pixel corresponds to 0.345 µm. Acquisition rates ranged from about 60 to 134 frames/s. Most recordings lasted about 15 s (median 14.9 s), and five long recordings were divided into 34 consecutive subrecordings. The recording library contains 2,759 analysis entries from 46 acquisition sessions.

#### Calcium imaging

Calcium imaging was performed with the Fluo-4 AM calcium indicator (Beyotime, #S1060). The medium was removed from the NRVMs, and the cells were washed three times with PBS. The probe was diluted to 2 µM in PBS, and the cells were incubated in it for 1 h at room temperature. The dye solution was then discarded, the cells were washed three times with PBS, and serum-free DMEM was added to maintain them. Fluorescence intensity was recorded on a fluorescence microscope.

#### Statistics

Paired comparisons of the same well before and after treatment used two-sided paired t tests. For the amplitude comparisons in Fig. 2C and Fig. S2, paired t tests were performed on log2-transformed amplitudes. Within each tool, comparisons between each drug concentration and the vehicle control used Dunnett’s test on the per-well ${log}_{2}$ fold change from before to after treatment, and Welch’s *t* tests with Holm correction across the five concentrations were used as a sensitivity analysis. Dose-response curves were fitted with a four-parameter log-logistic model (Fig. 2D and 2E). A well entered the analysis of a tool only when that tool detected at least one contraction in both the baseline and the treated recording. In the ISO series, this left five wells for CONTRACTIONWAVE at 100 nM and for Camystat at 10 nM and 100 nM. Agreement between readouts was assessed by Pearson correlation and by Bland-Altman analysis, which reports the bias and the 95% limits of agreement. Gap motion estimated by Cardiomyograph and by Farnebäck optical flow in the same recordings was compared with a two-sided Wilcoxon signed-rank test (Fig. S5C). Within-recording variability is reported as the coefficient of variation. Sample sizes and the units of n are given in the figure legends. Values are presented as mean ± SEM, and *P* < 0.05 was considered statistically significant.

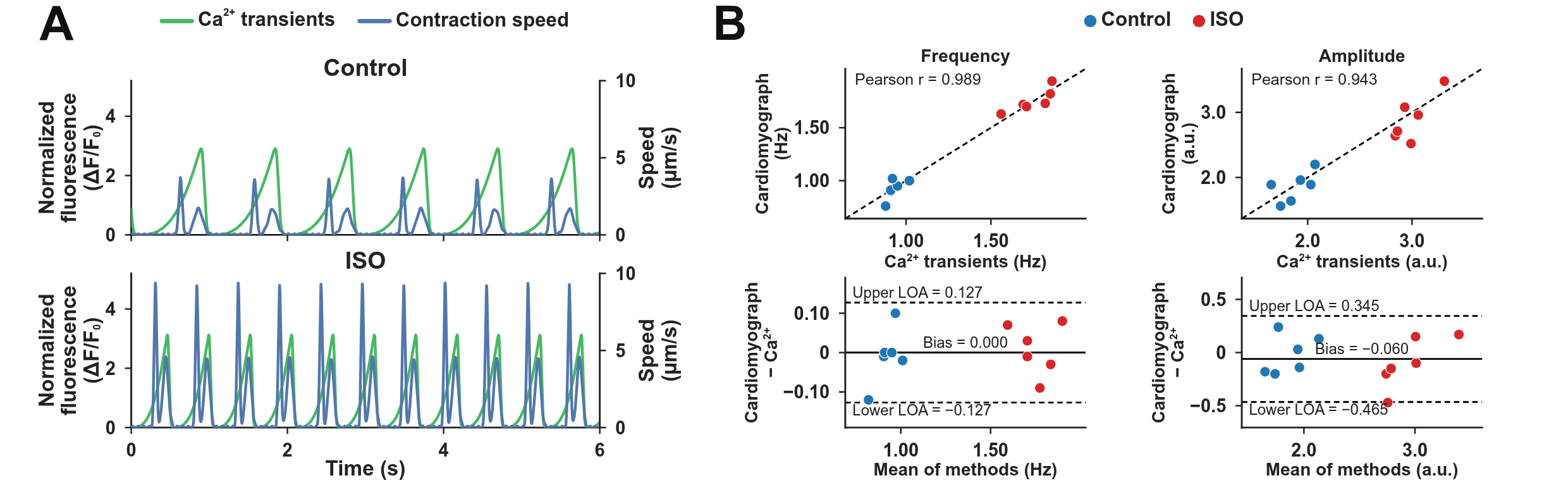

Figure S1. **Comparison of Cardiomyograph with Ca^2+^ imaging.**

**A,** Representative Cardiomyograph contraction speed and Ca^2+^ transients of neonatal rat ventricular myocytes (NRVMs) treated with vehicle (0.1% DMSO; Control) or isoprenaline (ISO, 1 µM).

**B,** Agreement between Cardiomyograph and Ca^2+^ transients in contraction frequency and amplitude across 12 recordings (Control 6, ISO 6). The top row shows the correlation, with the dashed line marking identity, and the bottom row shows the Bland-Altman analysis with 95% limits of agreement (LOA). a.u., arbitrary units; ΔF/F_0_, change in fluorescence relative to baseline fluorescence; DMSO, dimethyl sulfoxide; ISO, isoprenaline; LOA, limits of agreement; NRVMs, neonatal rat ventricular myocytes.

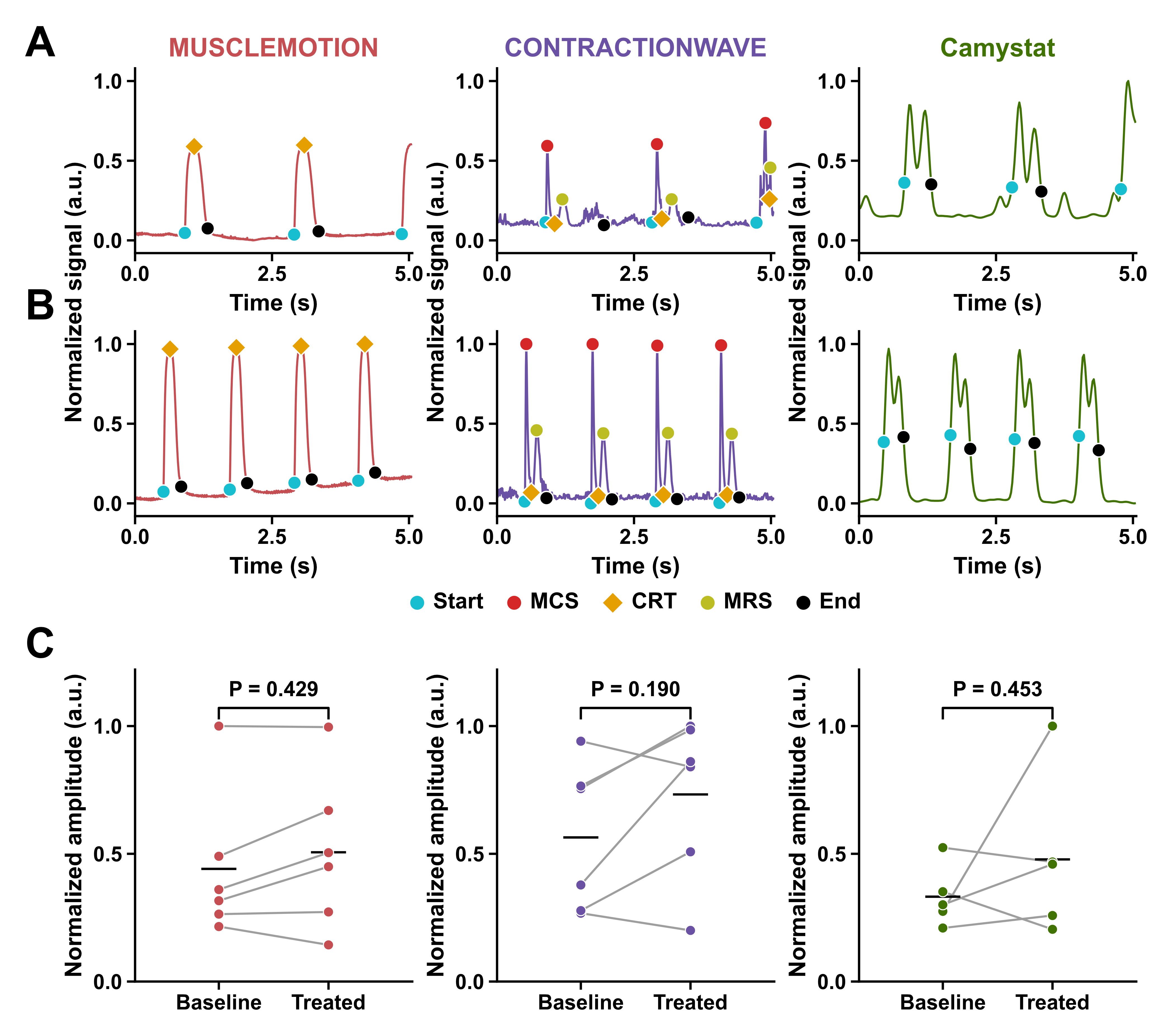

Figure S2. **Contraction amplitude in response to isoprenaline across three label-free contractility tools.**

**A and B,** Baseline (A) and treated (B) traces of one well exposed to 10 nM ISO. Within each tool, the two traces are normalized by a single factor. Symbols mark the start and end of each event detected by a tool and, where the tool reports them, the maximum contraction speed (MCS), the contraction-relaxation transition (CRT) and the maximum relaxation speed (MRS).

**C,** Amplitude of the baseline and treated recordings of the 10 nM wells (n = 6, 6, and 5 wells for MUSCLEMOTION, CONTRACTIONWAVE, and Camystat, respectively), with each panel divided by its largest value. Lines join the two recordings of one well, horizontal bars indicate group means, and P values are from two-sided paired t tests on log2 amplitude. a.u., arbitrary units; CRT, contraction-relaxation transition; ISO, isoprenaline; MCS, maximum contraction speed; MRS, maximum relaxation speed.

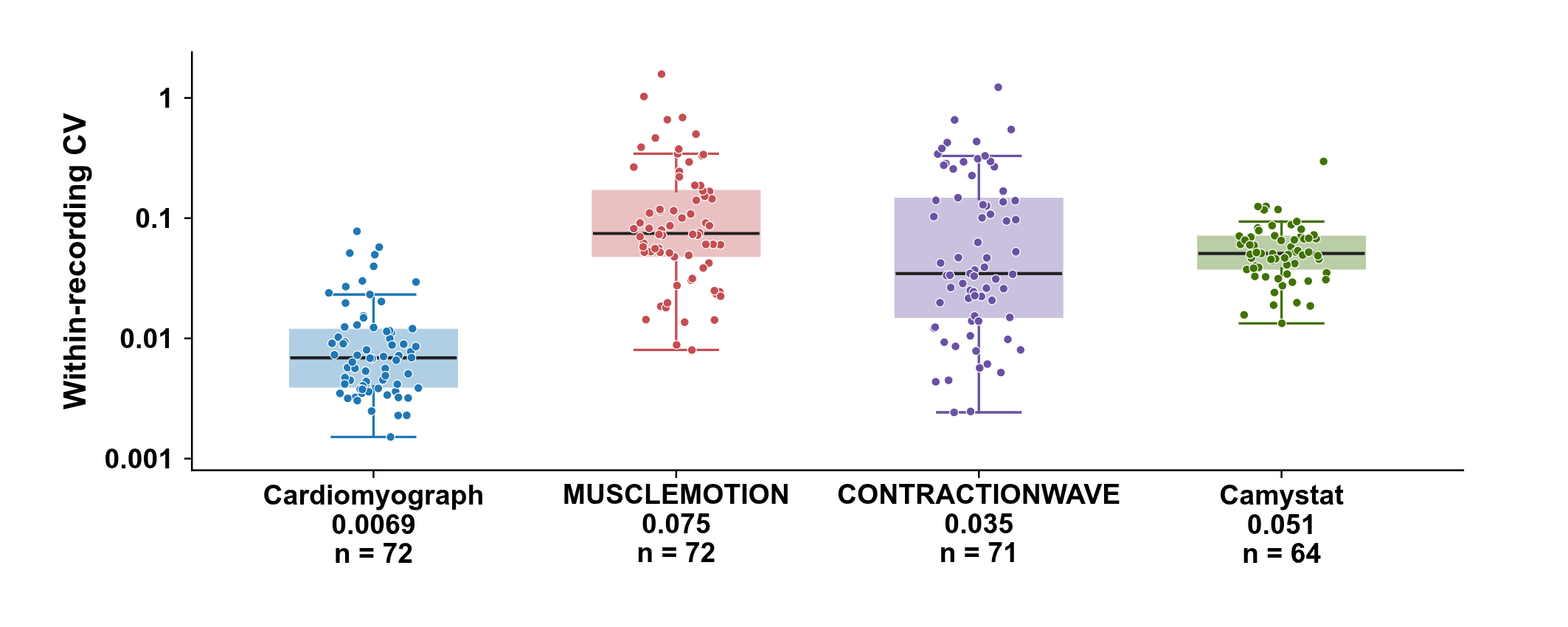

Figure S3. **Within-recording variability.**

Coefficient of variation (CV) of the per-contraction amplitude within each recording. Points are recordings, and numbers give the medians.

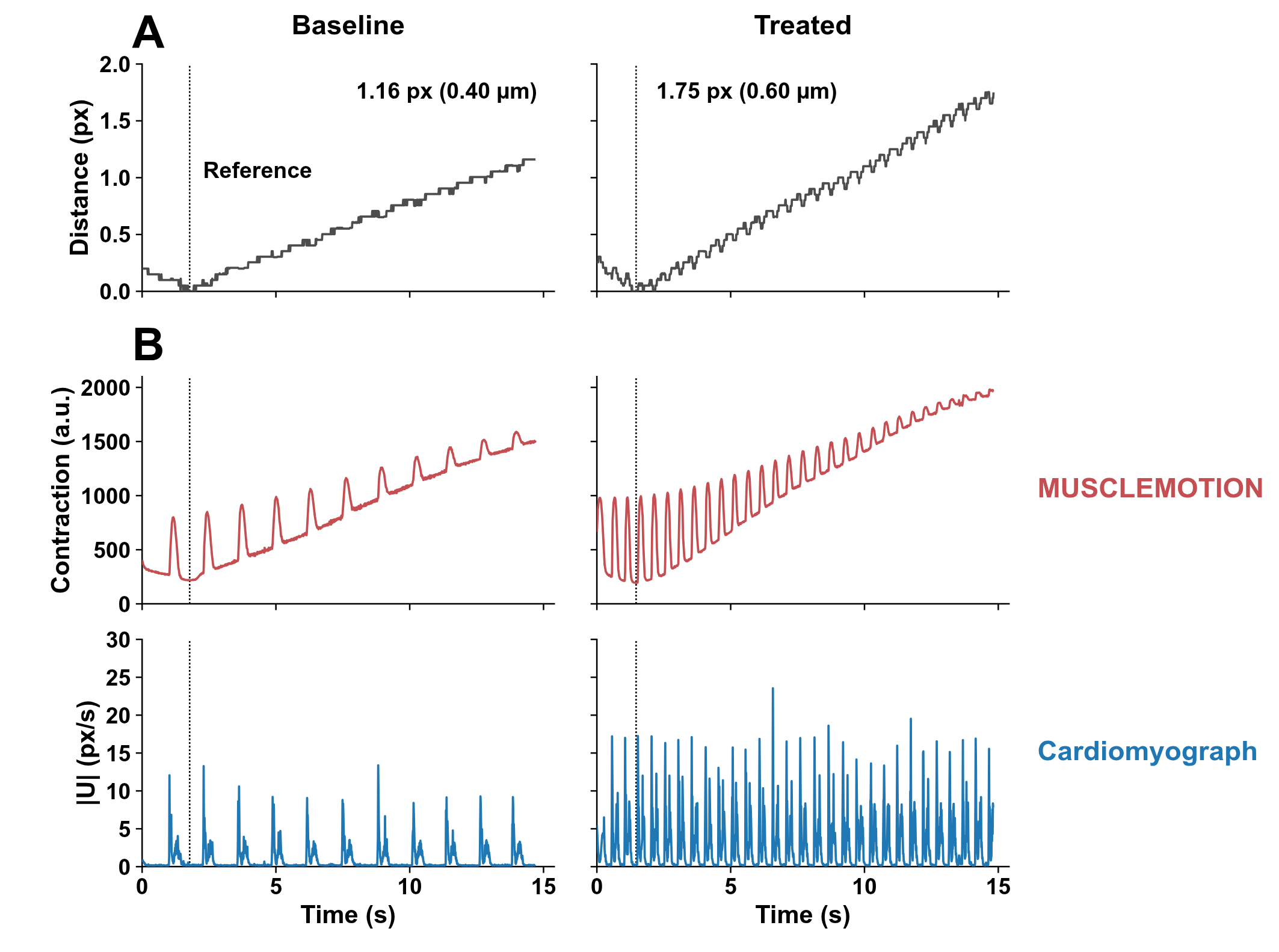

Figure S4. **Whole-field drift in the MUSCLEMOTION readout of one 10 nM ISO-treated pair.**

**A,** Distance of the whole field from its position in the MUSCLEMOTION reference frame (dotted lines).

**B,** MUSCLEMOTION contraction curve (top) and whole-field $|U|$ signal of Cardiomyograph (bottom) for the same recordings. a.u., arbitrary units; ISO, isoprenaline; $|U|$, motion-field magnitude.

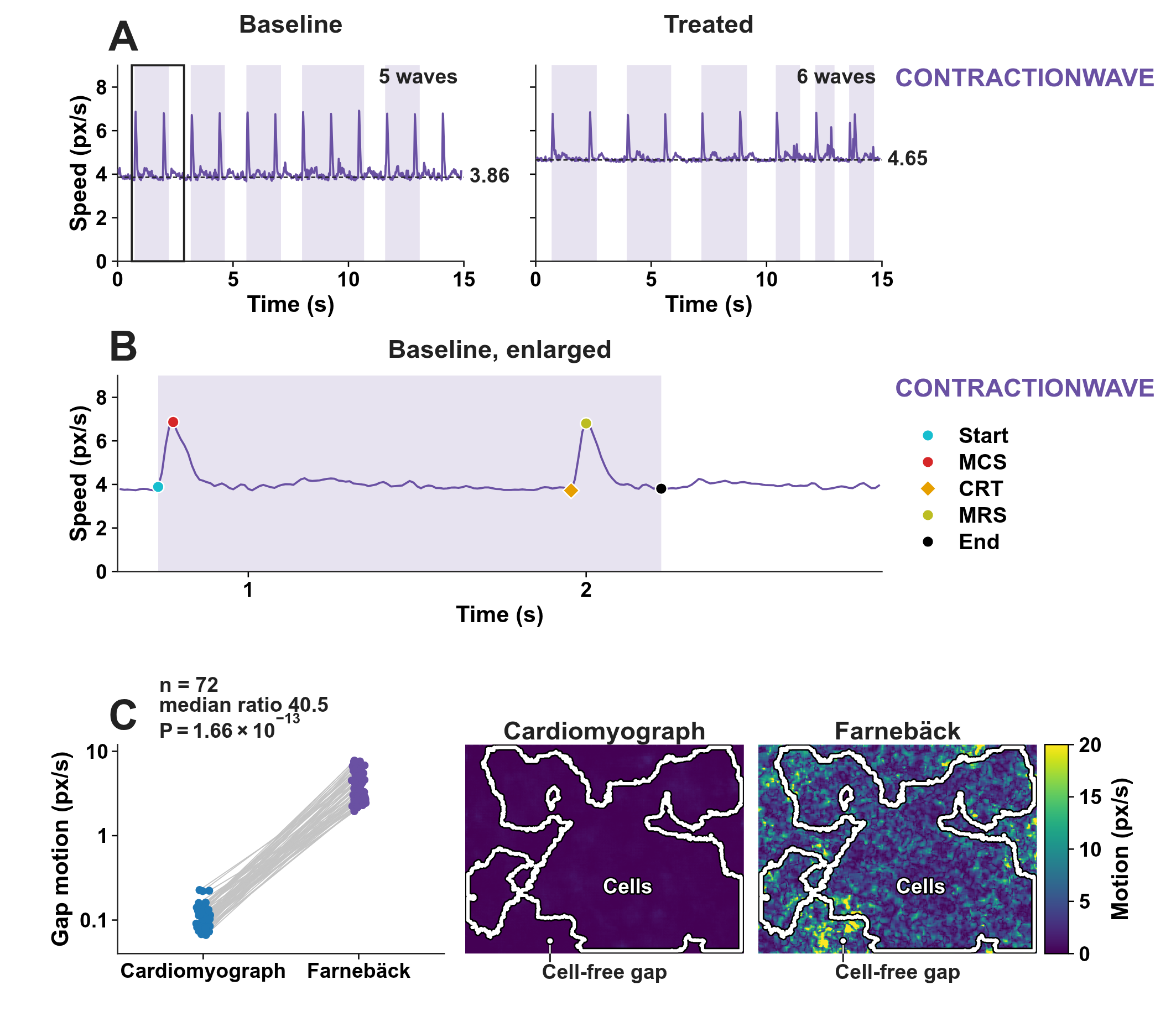

Figure S5. **Resting speed and detected waves in the CONTRACTIONWAVE readout of one 10 nM ISO-treated pair, and motion reported in cell-free gaps.**

**A,** CONTRACTIONWAVE speed. Dashed lines mark the resting level, and bands mark the detected waves.

**B,** Enlarged view of the boxed window in A.

**C,** Motion at rest in cell-free gaps, from Cardiomyograph and from the Farnebäck optical flow used by CONTRACTIONWAVE. Left, values for each recording (n = 72), joined by lines; two-sided Wilcoxon signed-rank test. Right, motion magnitude in one resting frame; the white line is the edge of the cell area. CRT, contraction-relaxation transition; ISO, isoprenaline; MCS, maximum contraction speed; MRS, maximum relaxation speed.

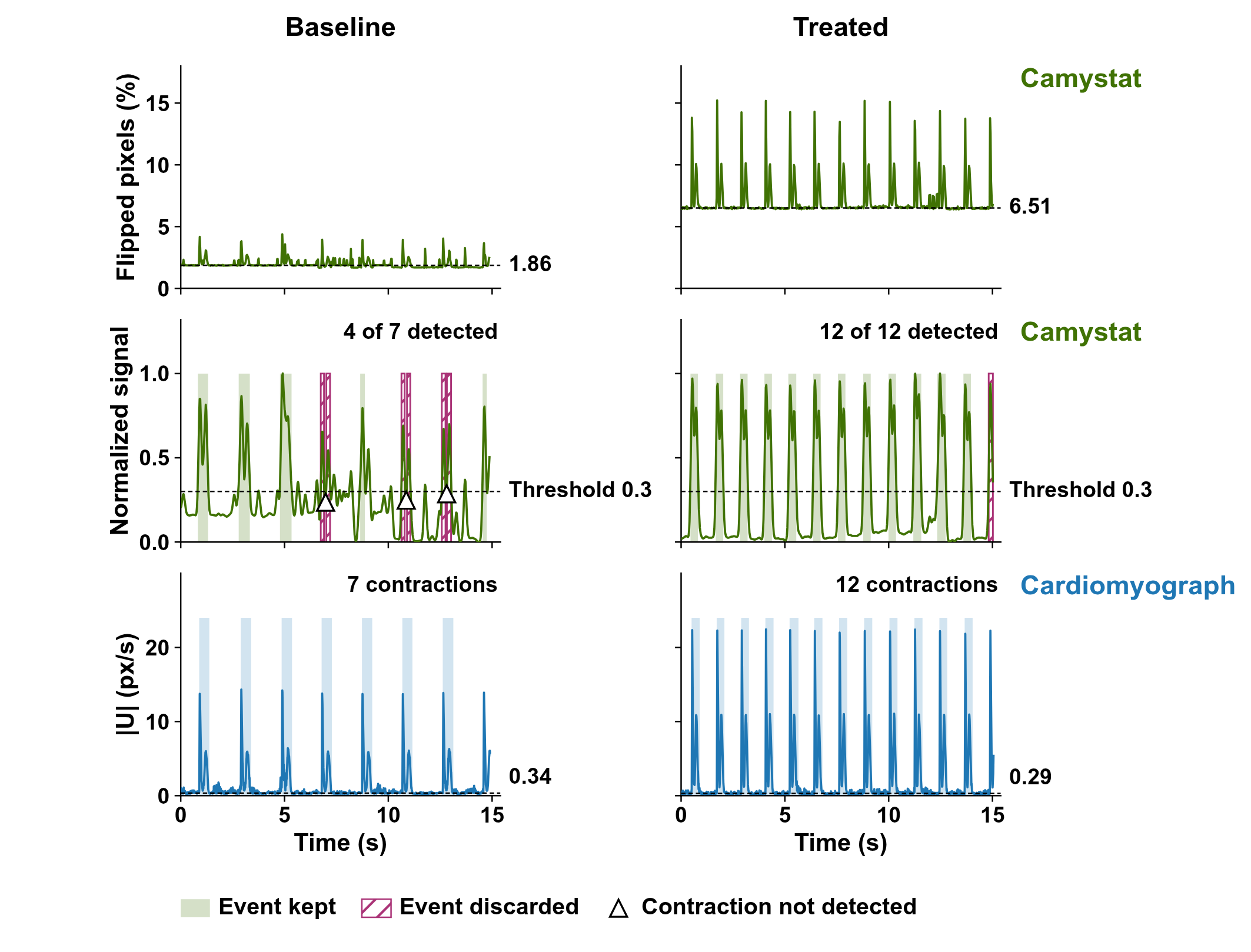

Figure S6. **Resting signal and missed contractions in the Camystat readout of one 10 nM ISO-treated pair.**

Top, percentage of flipped pixels reported by Camystat; dashed lines mark the resting level. Middle, normalized Camystat signal and its movement threshold (dashed lines). Filled bands mark kept events, hatched boxes mark discarded events, and triangles mark contractions without a matching event. Bottom, whole-field $|U|$ signal of Cardiomyograph; dashed lines mark the resting level, and bands mark its contractions. ISO, isoprenaline; $|U|$, motion-field magnitude.

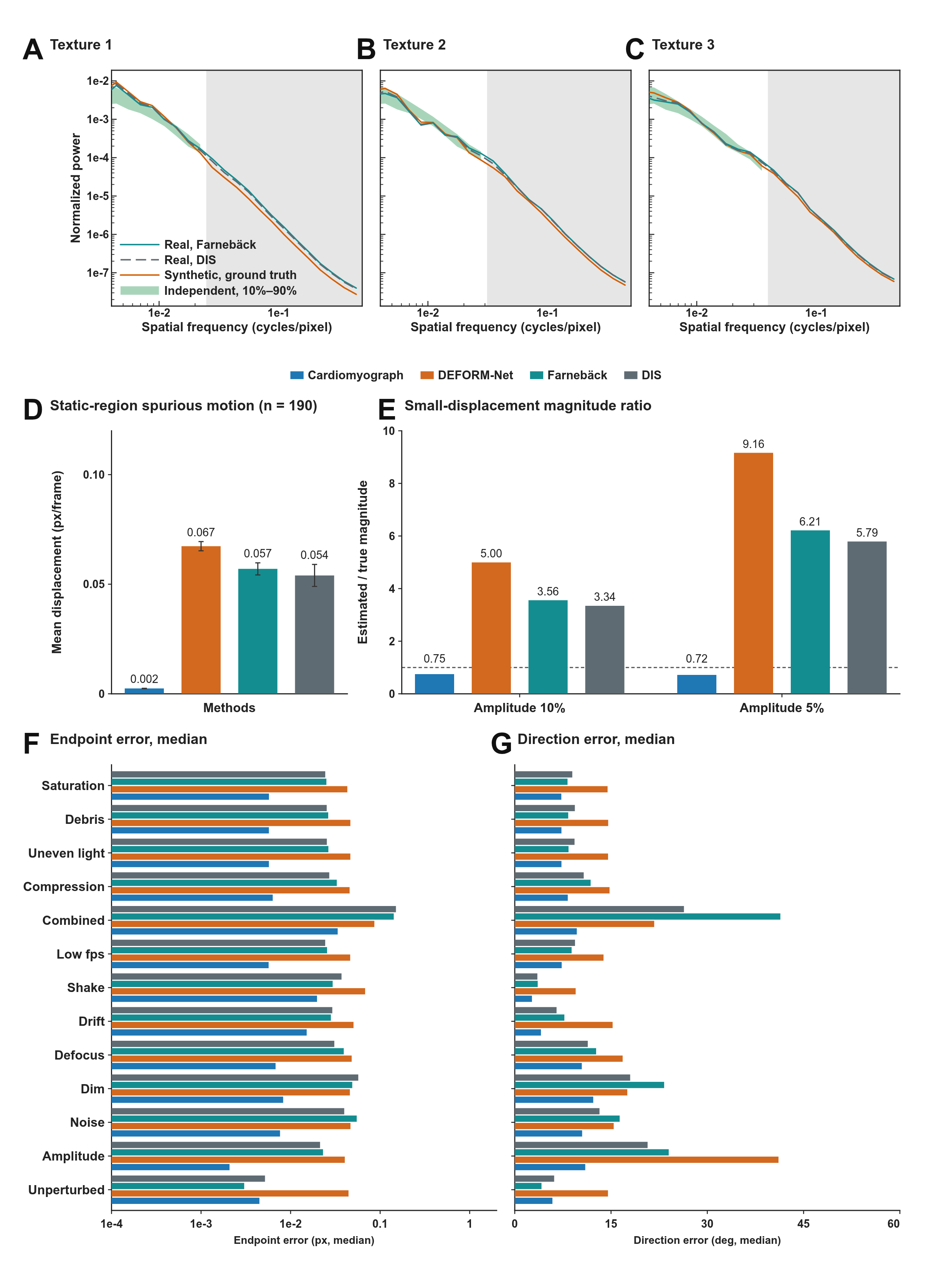

Figure S7. **Spatial power-spectral density of displacement in real and synthetic recordings, and motion-field estimation on synthetic recordings with known ground truth.**

**A-C,** Spatial power-spectral density (PSD) of displacement for Textures 1-3, estimated by optical flow on the source real recording (Real) and taken from the ground-truth displacement of the synthetic recording (Synthetic). The green band (Independent) is the 10th-90th percentile across 25 real recordings not used to generate synthetic recordings. Grey shading marks frequencies at which the ground-truth PSD is below 1% of its peak.

**D,** Spurious motion in static regions. Bars are means across recordings, and error bars indicate the standard error of the mean (SEM).

**E,** Estimated-to-true displacement-magnitude ratio at 10% and 5% of the unperturbed contraction amplitude. Bars are medians across recordings, and the dashed line marks a ratio of 1.

**F and G, Median endpoint error (F) and median direction error (G).** Bars are medians across the five textures. DIS, dense inverse search; fps, frames per second; PSD, power-spectral density; SEM, standard error of the mean.

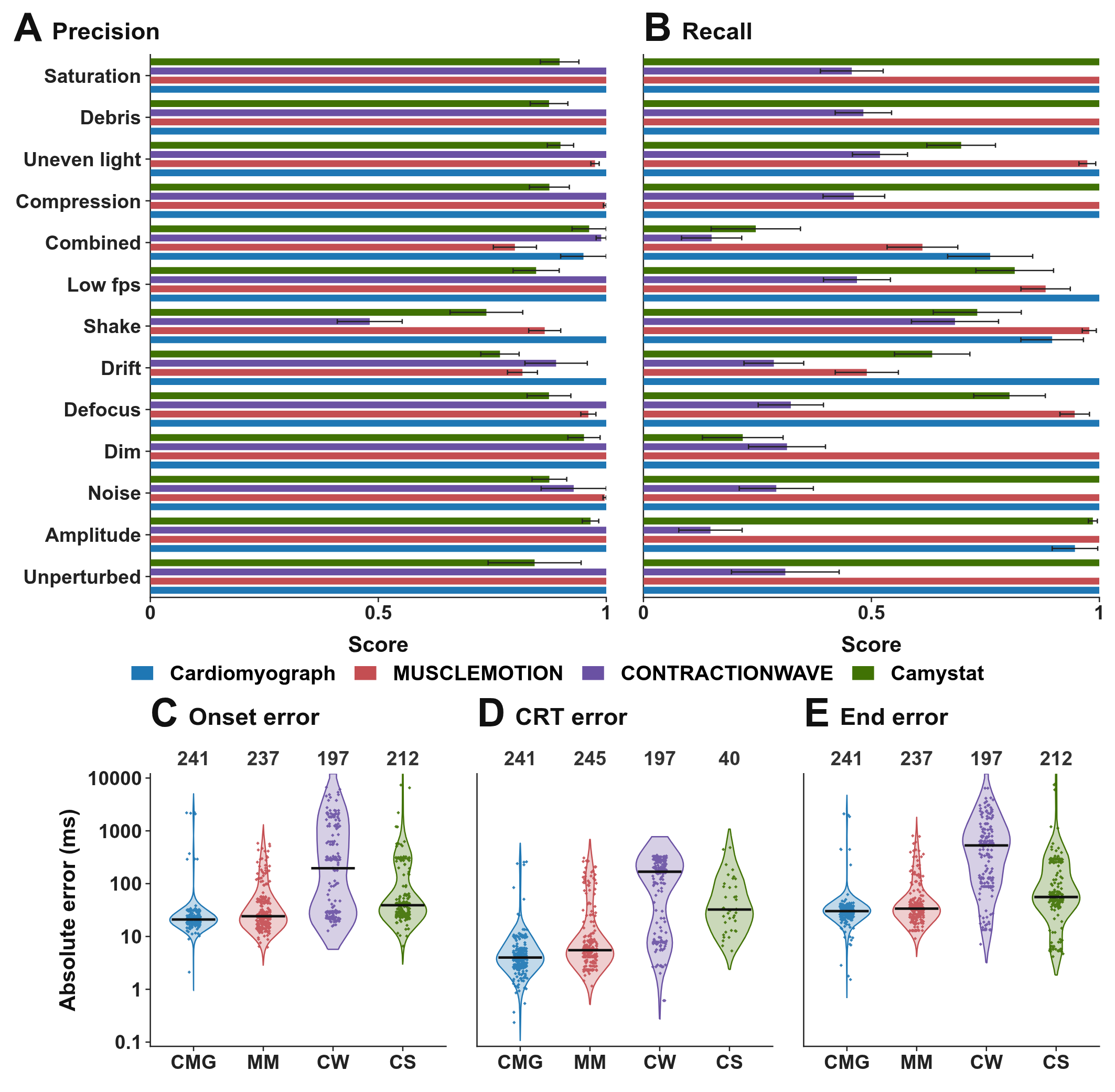

Figure S8. **Contraction detection and landmark timing on synthetic recordings with known ground truth.**

**A and B,** Precision (A) and recall (B) for the unperturbed recordings (n = 5) and for each of twelve perturbations (n = 20). Bars show means across recordings, and error bars indicate the standard error of the mean (SEM).

**C-E,** Absolute timing errors for onset (C), the contraction-relaxation transition (CRT, D), and the end of contraction (E). Points represent recording-level medians, black lines mark medians across recordings, and numbers above columns give the number of recordings. CMG, Cardiomyograph; CRT, contraction-relaxation transition; CS, Camystat; CW, CONTRACTIONWAVE; fps, frames per second; MM, MUSCLEMOTION; SEM, standard error of the mean.

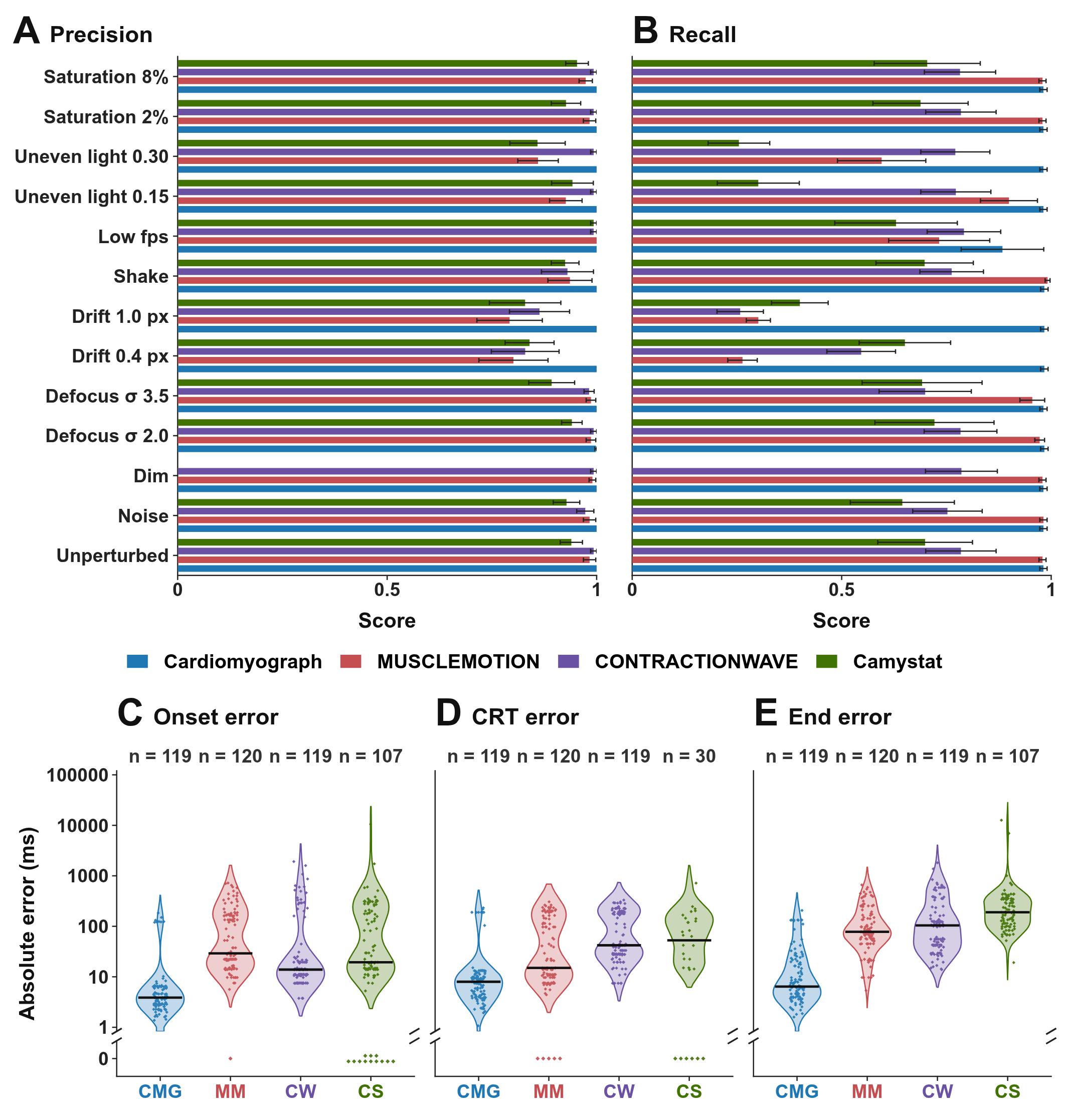

Figure S9. **Contraction detection and landmark timing on annotated real recordings with added perturbations.**

A and B, Precision (A) and recall (B) for 10 annotated recordings without perturbation and with each of 12 perturbations. Bars show means across recordings, and error bars indicate the standard error of the mean (SEM).

C-E, Absolute timing errors for onset (C), the contraction-relaxation transition (CRT, D), and the end of contraction (E) on the perturbed recordings. Points represent recording-level medians, black lines mark medians across recordings, and numbers above columns give the number of recordings. CMG, Cardiomyograph; CRT, contraction-relaxation transition; CS, Camystat; CW, CONTRACTIONWAVE; fps, frames per second; MM, MUSCLEMOTION; SEM, standard error of the mean.

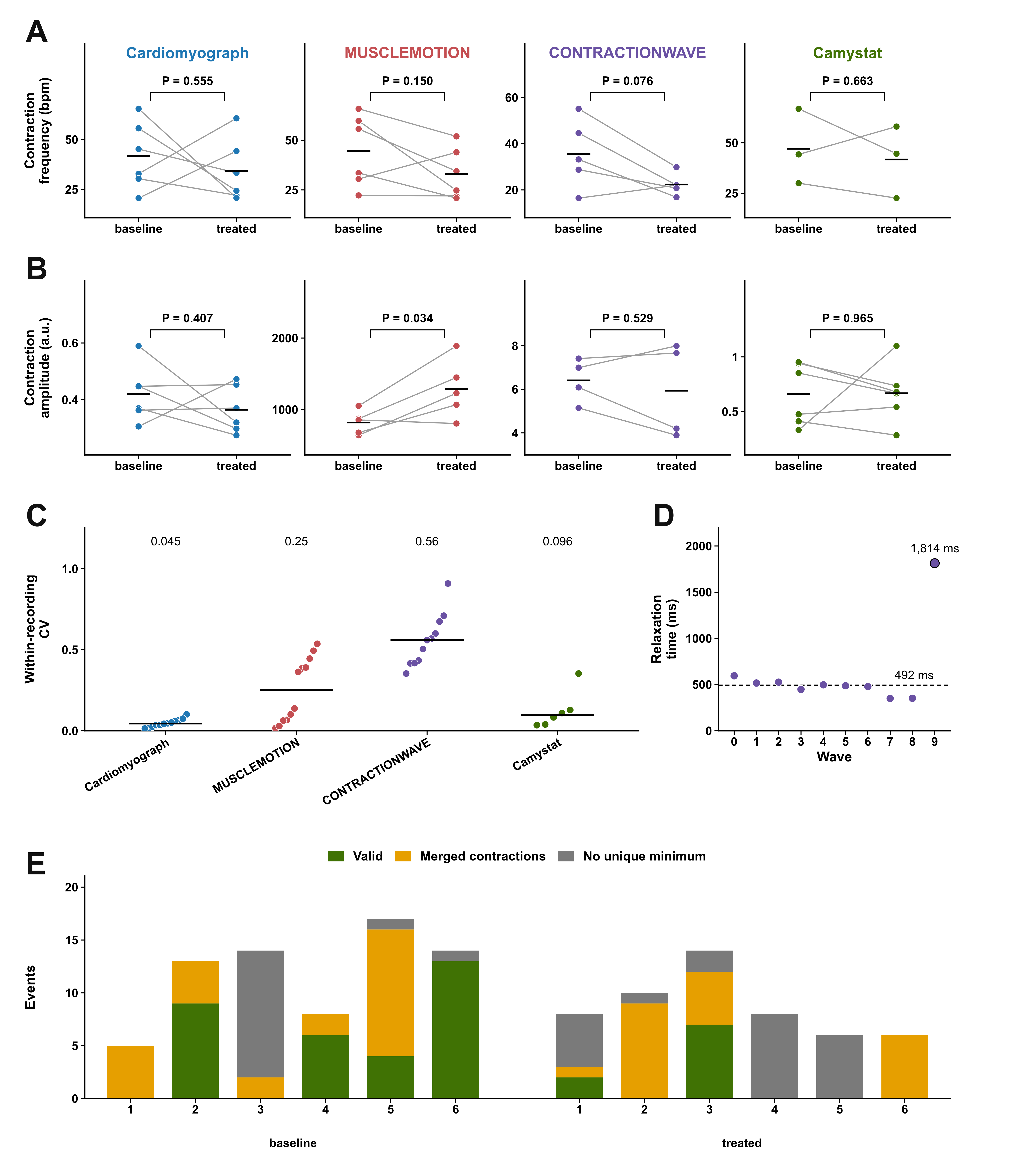

Figure S10. **Paired contractile readouts and within-recording variation before and after E-4031 at 50 nM.**

**A and B,** Paired contraction frequency (A) and contraction amplitude (B). Each dot is the median value for one recording. Lines connect paired recordings, and horizontal bars indicate group means. Only baseline-treated pairs with available values are shown (A: n = 6, 6, 5, and 3 pairs; B: n = 6, 5, 4, and 6 pairs for Cardiomyograph, MUSCLEMOTION, CONTRACTIONWAVE, and Camystat, respectively). P values are from two-sided paired t tests.

**C,** Within-recording coefficient of variation of relaxation time. Each point represents one recording, and the black horizontal bar indicates the median for that tool (n = 12, 12, 11, and 6 recordings for Cardiomyograph, MUSCLEMOTION, CONTRACTIONWAVE, and Camystat, respectively).

**D,** Relaxation times of the ten CONTRACTIONWAVE-detected waves in one recording. The dashed line indicates the median, and the outlined point marks the longest relaxation time.

**E,** Camystat event counts for six baseline and six treated recordings. a.u., arbitrary units; bpm, beats per minute; CV, coefficient of variation.

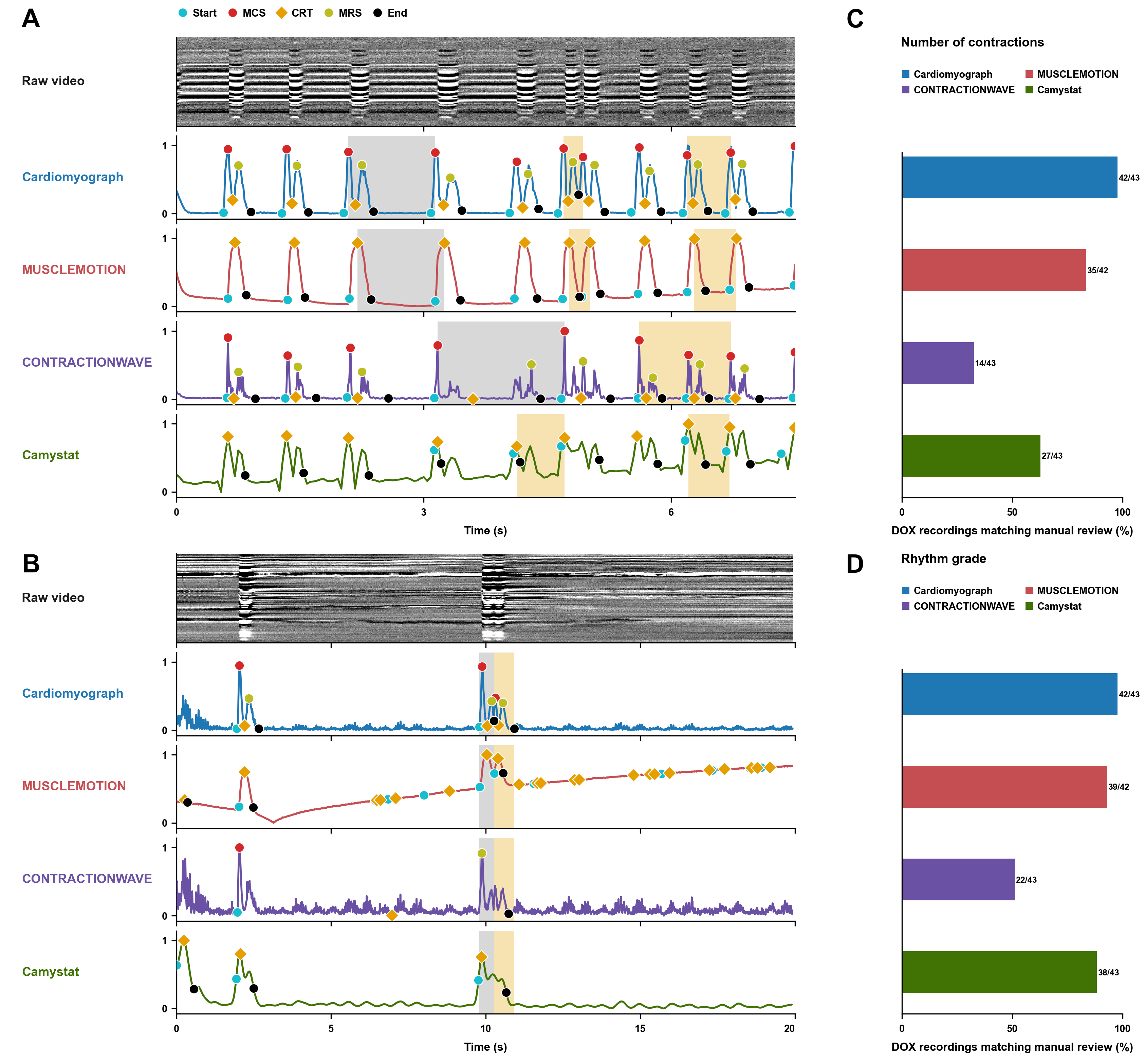

Figure S11. **Irregular contraction rhythms** after **oxygen-glucose deprivation (OGD) or doxorubicin** (DOX), and agreement of each tool with manual review in the DOX experiment.

A and B, An OGD-treated recording (A) and a recording of NRVMs after 20 h of exposure to 0.5 µM DOX (B). The top row shows a space-time slice of the raw video, and the four rows below show the contraction signals from Cardiomyograph, MUSCLEMOTION, CONTRACTIONWAVE, and Camystat. In A, coloured bands mark intervals that deviate from the local median contraction interval. In B, the grey band marks the first cycle after a long pause, and the orange band marks the cycle that follows it.

C and D, Percentage of DOX recordings in which each tool detected the same number of contractions (C) and gave the same rhythm grade (D) as the manual review. Fractions give the number of matching recordings over the number analysed.

CRT, contraction-relaxation transition; DOX, doxorubicin; MCS, maximum contraction speed; MRS, maximum relaxation speed; NRVMs, neonatal rat ventricular myocytes; OGD, oxygen-glucose deprivation.

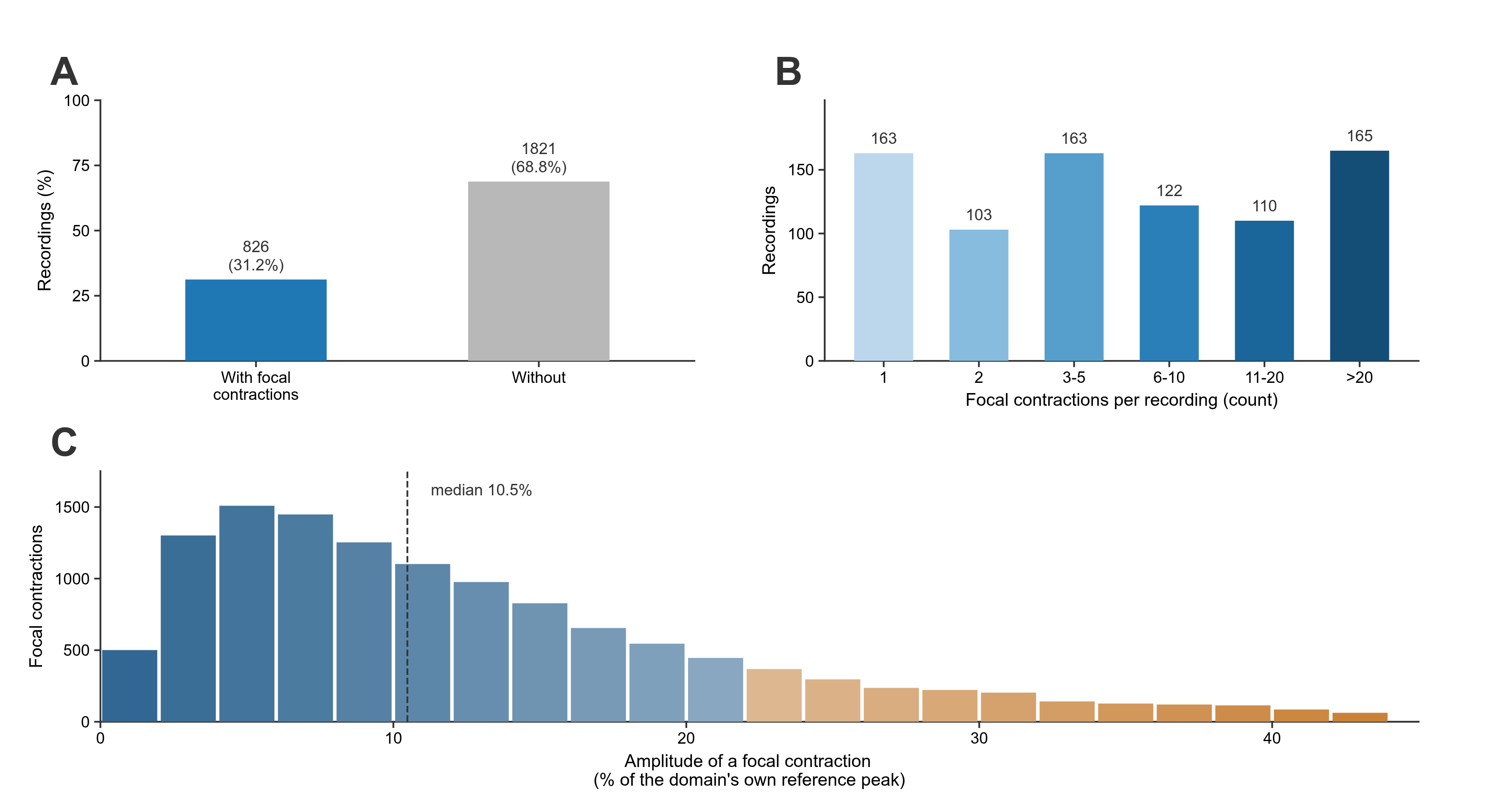

**Figure S12. Prevalence and amplitude of focal contractions in all recordings with contractions.**

**A,** Recordings with and without focal contractions among the 2,647 recordings with contractions.

**B,** Number of focal contractions per recording, among the 826 recordings with at least one focal contraction.

**C,** Amplitude of each focal contraction as a percentage of its domain's reference peak (n = 12,580). The dashed line marks the median, and the 45% threshold sets the right end of the distribution.

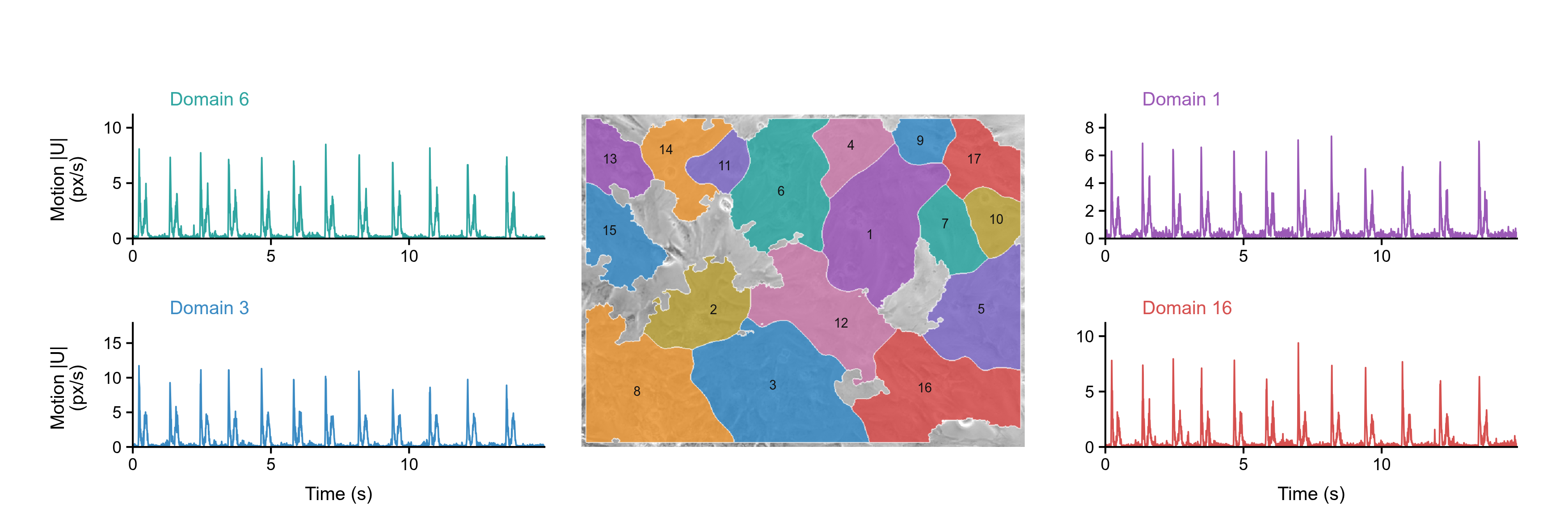

Figure S13. **Domain-level analysis of one recording.**

Centre, the field of view overlaid with the domain segmentation; each domain has one colour and one number. Left and right, the motion trace (|U|) of four domains, each drawn in the colour of its domain. Each vertical scale is set from the peak of its own trace. px/s, analysis pixels per second; $|U|$, motion-field magnitude.

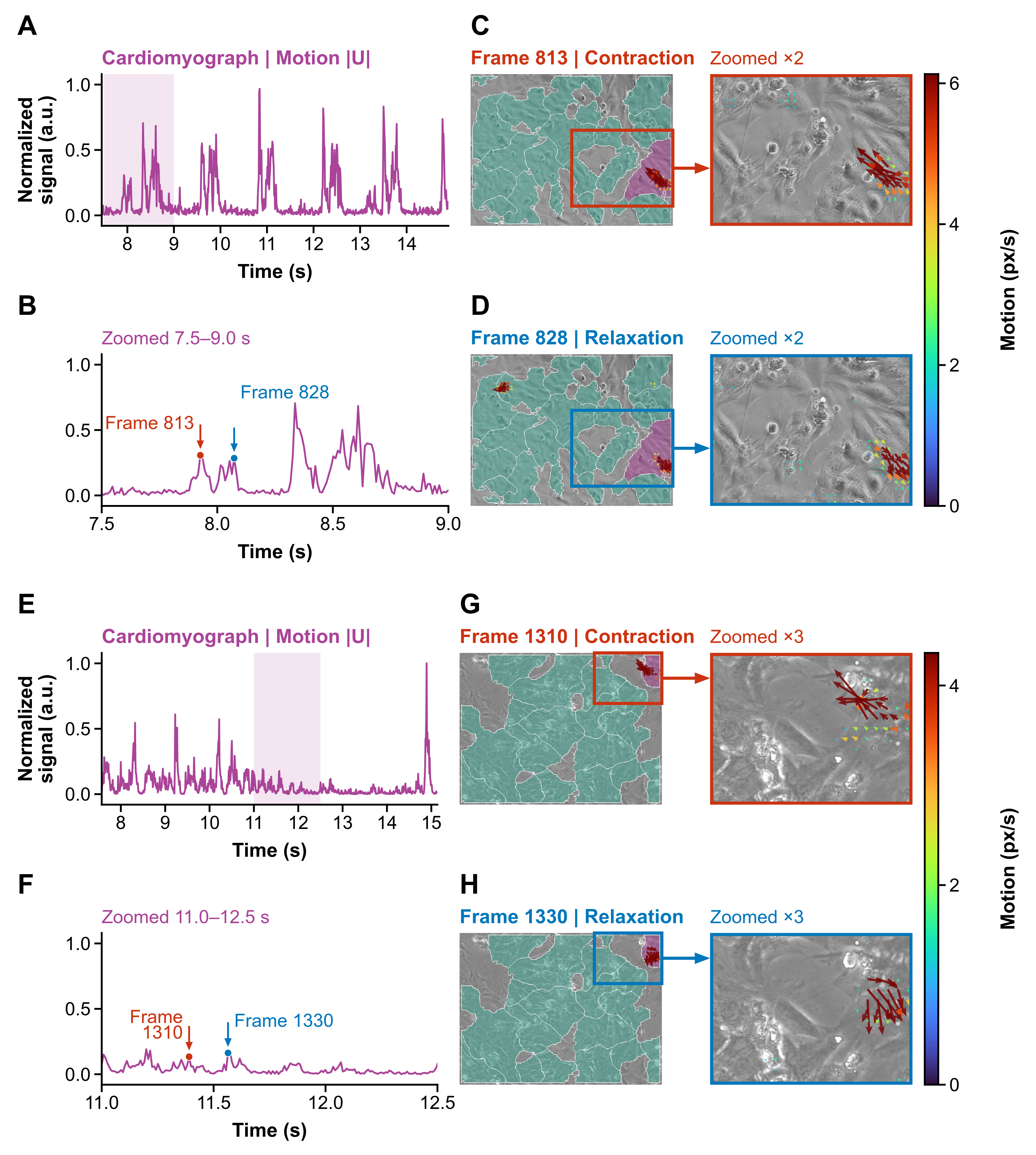

Figure S14. **Focal events in two further recordings.**

**A,** Cardiomyograph output for the recording with 23 domains. The motion signal |U| of the focal domain is normalized to 0-1, and the shaded band marks the window shown in B.

**B,** The 7.5-9.0 s window. The red arrow marks frame 813, during a focal contraction, and the blue arrow marks frame 828, during the following relaxation.

**C and D,** Frames 813 (C) and 828 (D). The focal domain is shown in purple and the other domains in teal, and the boxed region is enlarged 2×. Arrows give the local direction of motion, and colour gives the magnitude in px/s.

**E-H,** The recording with 17 domains, shown as in A-D, with frames 1310 and 1330 marked in the 11.0-12.5 s window and the boxed region enlarged 3×. a.u., arbitrary units; $|U|$, motion-field magnitude.

### Supplementary tables

**Table S1.** Contraction-level parameters, computed for each contraction of each domain.

| **No.** | **Parameter** | **Unit** | **Definition** |
| --- | --- | --- | --- |
| 1 | Onset-to-end duration | ms | Time from onset $\tau_{0}$ to end $\tau_{4}$. |
| 2 | 50% displacement recovery time | ms | Time from the displacement peak until the displacement first falls halfway from the peak to its value at End (ms). |
| 3 | Time to peak displacement | ms | Time from contraction onset to the peak of the onset-subtracted displacement trace (ms). |
| 4 | 90% speed recovery after MRS | s | Time from MRS until speed has fallen 90% of the way from the MRS value to the contraction-end value; if that crossing is missing, the whole MRS-to-end interval (s). |
| 5 | Displacement width at half-maximum | ms | Time between the rising and the falling crossing of the level used for the 50% displacement recovery time (ms). |
| 6 | Relaxation time constant | ms | Exponential time constant of the displacement decay, fitted from the 50% recovery point to End; kept only if fit $R^{2}$ is at least 0.80, the solution is below the upper fitting bound, and it is shorter than 10 times the fitted interval. |
| 7 | 50% speed recovery after MRS | s | Time from MRS until speed has fallen halfway from the MRS value to the contraction-end value; if that crossing is missing, the whole MRS-to-end interval (s). |
| 8 | Speed width at half-maximum | s | Duration from the first to the last sample at or above the minimum of the onset, $t_{2}$, and end amplitudes plus half the rise to the contraction maximum; 0 s when fewer than two samples qualify. |
| 9 | Extra speed-peak count | count | Count of local speed peaks at least $max(0.01,0.2\text{ times the main peak},0.3\text{ standard deviations (SD)})$, excluding MCS and MRS within one frame (count). |
| 10 | Time to 50% of MCS | s | Time from onset until speed first reaches halfway from the onset value to the MCS value; 0 s if that crossing is absent. |
| 11 | Time to 90% of MCS | s | Time from onset until speed first reaches 90% of the rise from the onset value to the MCS value; 0 s if that crossing is absent. |
| 12 | Contraction synchrony index | 1 | Variance of the mean z-scored speed traces of the member domains of the synchronous domain, taken from their median Start to their median CRT, divided by the mean variance of those traces, clipped to 0-1 (1). |
| 13 | Speed at MRS | analysis-px/s | $\parallel U\parallel(t_{\mathrm{MRS}})$. |
| 14 | Peak mode displacement | mode-score×s per pixel | Maximum of the displacement between $\tau_{0}$ and $\tau_{4}$, divided by the number of pixels in the domain. |
| 15 | Relaxation displacement integral | mode-score×s per pixel | Time integral of $S$, relative to its value at $\tau_{0}$, over the closed interval from $\tau_{2}$ to $\tau_{4}$, divided by the number of pixels in the domain. |
| 16 | Contraction-phase duration | ms | Time from $\tau_{0}$ to $\tau_{2}$. |
| 17 | Relaxation duration | ms | Time from $\tau_{2}$ to $\tau_{4}$. |
| 18 | Time to MCS | ms | Time from $t_{0}$ to MCS. |
| 19 | Time from MRS to end | ms | Time from MRS to $t_{4}$. |
| 20 | CRT-to-MRS time | ms | Time from CRT to MRS. |
| 21 | 90% displacement recovery time | ms | Time from the displacement peak until the displacement first falls 90% of the way from the peak to its value at End (ms). |
| 22 | Speed at MCS | analysis-px/s | $\parallel U\parallel(t_{\mathrm{MCS}})$. |
| 23 | Speed at CRT | analysis-px/s | $\parallel U\parallel(t_{2})$. |
| 24 | Contraction speed integral | analysis-px | Trapezoidal integral over time, from $\tau_{0}$ to $\tau_{2}$, of the part of $\parallel U\parallel$ above the lower of its values at $\tau_{0}$ and $\tau_{2}$. This integrates the speed trace and is distinct from the contraction displacement integral. |
| 25 | Contraction displacement integral | mode-score×s per pixel | Time integral of $S$, relative to its value at $\tau_{0}$, over the closed interval from $\tau_{0}$ to $\tau_{2}$, divided by the number of pixels in the domain. This integrates the mode score and is distinct from the contraction speed integral. |
| 26 | Late-relaxation slope | analysis-px/s^2^ | Change in speed from the midpoint of MRS-to-end to the contraction end, divided by that time span. |
| 27 | Instantaneous rate | min^−1^ | Rate from the interval between this onset and the next onset. |
| 28 | Baseline recovery error | analysis-px/s | Absolute difference of $\parallel U\parallel$ between $t_{4}$ and $t_{0}$. |
| 29 | Relaxation-fit $R^{2}$ | 1 | $R^{2}$ of the exponential fit to the displacement decay from the 50% recovery point to End. |
| 30 | Active-domain fraction | 1 | Fraction of the area of the synchronous domain taken by member domains that have their own landmarks for that contraction. |
| 31 | Active-domain count | count | Count of member domains of the synchronous domain that have their own landmarks for that contraction. |

**Table S2.** Domain-level parameters, computed for each domain.

| **No.** | **Parameter** | **Unit** | **Definition** |
| --- | --- | --- | --- |
| 1 | Robust CV of contraction-phase duration | 1 | Median of 20 deterministic eight-event subsamples of the interquartile range divided by the absolute median of contraction duration. |
| 2 | Classic CV of peak speed | 1 | Median of 20 deterministic eight-event subsamples of the sample standard deviation divided by the absolute mean of absolute MCS amplitude. |
| 3 | Classic CV of contraction-phase duration | 1 | Median of 20 deterministic eight-event subsamples of the sample standard deviation divided by the absolute mean of contraction duration. |
| 4 | rIQR of contraction-phase duration | 1 | IQR of contraction duration divided by the absolute median. |
| 5 | rIQR of relaxation duration | 1 | IQR of relaxation duration divided by the absolute median. |
| 6 | rIQR of time to MCS | 1 | IQR of time to MCS divided by the absolute median. |
| 7 | rIQR of time to peak displacement | 1 | IQR of time to peak displacement divided by the absolute median. |
| 8 | rIQR of 50% displacement recovery time | 1 | IQR of 50% displacement recovery time divided by the absolute median. |
| 9 | rIQR of 90% displacement recovery time | 1 | IQR of 90% displacement recovery time divided by the absolute median. |
| 10 | Contraction-duration span | ms | 90th minus 10th percentile of contraction duration. |
| 11 | Contraction-duration trend | ms/contraction | Least-squares slope of contraction duration against contraction order. |
| 12 | IQR of the onset-to-MCS phase | cycle fraction | IQR across contractions of (MCS minus onset) divided by the full-contraction duration. |
| 13 | IQR of the MCS-to-CRT phase | cycle fraction | IQR across contractions of ($t_{2}$ minus MCS) divided by the full-contraction duration. |
| 14 | IQR of the CRT-to-MRS phase | cycle fraction | IQR across contractions of (MRS minus $t_{2}$) divided by the full-contraction duration. |
| 15 | IQR of the MRS-to-end phase | cycle fraction | IQR across contractions of (end minus MRS) divided by the full-contraction duration. |
| 16 | Median shape correlation | correlation | Median Pearson correlation of normalized contraction waveforms with their mean template. |
| 17 | Shape-correlation IQR | correlation | IQR of Pearson correlations of normalized contraction waveforms with their mean template. |
| 18 | Median waveform concordance | 1 | Median concordance correlation of raw waveforms with their mean template. |
| 19 | Waveform-concordance IQR | 1 | IQR of concordance correlations of raw waveforms with their mean template. |
| 20 | Deformed-contraction fraction | 1 | Fraction of contractions whose shape correlation is below 0.95. |
| 21 | Strength trend | 1/contraction | Ordinary-least-squares slope of the median-normalized warped-waveform peak against contraction order. |
| 22 | Strength residual IQR | 1 | Interquartile range of residuals after regressing the median-normalized warped-waveform peak on contraction order and even-odd alternation. |
| 23 | Robust CV of peak speed | 1 | Median of 20 deterministic eight-event subsamples of the interquartile range divided by the absolute median of absolute MCS amplitude. |
| 24 | Peak-speed span | analysis-px/s | 90th minus 10th percentile of absolute MCS amplitude. |
| 25 | Peak-speed half difference | analysis-px/s | Median absolute MCS amplitude in the second half of contractions minus that in the first half. |
| 26 | Median inter-contraction interval | s | Median interval between successive contraction onsets in the synchronous domain to which the domain belongs, with contractions truncated by the start or end of the recording excluded. |
| 27 | Inter-contraction-interval SD^a^ | s | Sample standard deviation, divisor n-1, of the inter-contraction intervals of the synchronous domain; requires at least 2 intervals. |
| 28 | Interval lag-1 correlation | correlation | Correlation between successive inter-contraction intervals; requires at least 8 intervals. |
| 29 | Interval-next-peak correlation | correlation | Correlation between an inter-contraction interval and the next absolute MCS amplitude; requires at least 7 intervals. |
| 30 | Direction dispersion | 1 | One minus the length of the summed contraction-flow vectors, divided by the sum of their magnitudes. |
| 31 | Centripetal index | 1 | Inward radial component of gated contraction flow, divided by the sum of flow magnitudes. |
| 32 | Top-five singular-energy fractions | 1 | The first five eigenvalue fractions of the domain residual flow (1). |
| 33 | First-mode energy fraction | 1 | Fraction of residual-flow energy in the first singular mode (1). |
| 34 | rIQR of the inter-contraction interval | 1 | IQR of inter-contraction intervals divided by the absolute median. |
| 35 | Strength alternation | 1 | Lag-1 correlation of the median-normalized warped-waveform peak after removing the trend versus contraction order, computed on pairs of consecutive contractions; requires at least 7 such pairs. |

^a^Interval CV is the SD in row 27 divided by the mean of the same intervals, computed when at least 2 intervals are available.
